# Deep learning model of somatic hypermutation reveals importance of sequence context beyond targeting of AID and Polη hotspots

**DOI:** 10.1101/2021.08.03.453264

**Authors:** Catherine Tang, Artem Krantsevich, Thomas MacCarthy

## Abstract

B-cells undergo somatic hypermutation (SHM) of the Immunoglobulin (Ig) variable region to generate high-affinity antibodies. SHM relies on the activity of activation-induced deaminase (AID), which mutates C>U preferentially targeting WR<u>C</u> (W=A/T, R=A/G) hotspots. Downstream mutations at W<u>A</u> Polymerase η hotspots contribute further mutations. Computational models of SHM can describe the probability of mutations essential for vaccine responses. Previous studies using short subsequences (*k*-mers) failed to explain divergent mutability for the same *k*-mer. We developed the DeepSHM (Deep learning on SHM) model using *k*-mers of size 5-21, improving accuracy over previous models. Interpretation of DeepSHM identified an extended DWR<u>C</u>T (D=A/G/T) motif with particularly high mutability. Increased mutability was further associated with lower surrounding G content. Our model also discovered a conserved AGYC<u>T</u>GGGGG (Y=C/T) motif within FW1 of IGHV3 family genes with unusually high T>G substitution rates. Thus, a wider sequence context increases predictive power and identifies novel features that drive mutational targeting.

## Introduction

Upon encountering antigen, germinal center (GC) B cells undergo several programmed mutational events in secondary lymphoid organs to mount an effective humoral immune response. Somatic hypermutation (SHM) takes place in the GC dark zone whereby mostly point mutations are introduced into the Immunoglobulin (Ig) variable (V) region. Selection for mutations leading to higher binding B cell receptors to cognate antigen occurs in the GC light zone, thus, producing a diverse repertoire of high-affinity antibodies (Methot and Di Noia, 2017; Pilzecker and Jacobs, 2019; Rajewsky, 1996). The mutagenic enzyme, activation-induced deaminase (AID), initiates SHM (Muramatsu et al., 2000) by converting cytosine (C) to uracil (U) in single-stranded DNA (ssDNA), resulting in a U:G (guanine) mismatch (Bransteitter et al., 2003). AID displays preferential targeting at WR<u>C</u>/<u>G</u>YW “hotspot” motifs (where W=A/T, R=A/G, Y=C/T, and the underlined base indicates the mutated base in the top and bottom strand, respectively), whereas SY<u>C</u>/<u>G</u>RS “coldspots” (S=C/G) are significantly less targeted (Pham et al., 2003; Rogozin and Diaz, 2004; Rogozin and Kolchanov, 1992; Yu et al., 2004). If left unrecognized, U mismatches will act as a template T and be replicated over (Pilzecker and Jacobs, 2019). The resulting C>T transition mutation is commonly referred to as the DNA “footprint” of AID (Liu et al., 2008). Downstream DNA repair further contributes to antibody diversity that is mediated by low-fidelity polymerases. During non-canonical base-excision repair (ncBER), the U:G mismatch is recognized and excised by uracil-DNA glycosylase (UNG), resulting in an abasic site (Rada et al., 2004). Repair of these abasic sites by REV1 can cause both transition and transversion mutations at C:G base-pairs (Jansen et al., 2006). In the case of non-canonical mismatch repair (ncMMR), the U:G mismatch is recognized by the MSH2/MSH6 heterodimer. Next, EXO1 exonuclease is recruited to create a patch of ssDNA, which then allows error-prone polymerases, particularly Polymerase eta (Polη), to resynthesize. Polη is known to create mutations at neighboring adenine (A) and thymine (T) sites of the initial AID-induced lesion, most notably at W<u>A</u>/<u>T</u>W hotspot motifs (Matsuda et al., 2001; Mayorov et al., 2005).

Several computational models have been developed for the SHM process and intrinsic biases exhibited by key proteins such as AID and Polη. These models have mainly utilized *k*-mer subsequences, where *k* is a specified integer length, ranging between 3- to 7-mers (Cui et al., 2016; Elhanati et al., 2015; Shapiro et al., 1999; Shapiro et al., 2002; Yaari et al., 2013). Two of these models (Cui et al., 2016; Yaari et al., 2013) are widely used and have leveraged 5-mer motifs to capture the dependency of the local surrounding sequence for the middle nucleotide to mutate, while simultaneously bypassing any influence of selection. The first of these targeting models (“S5F”) evaluates all possible 5-mers and synonymous (silent) mutations derived from functionally rearranged, or productive, VDJ coding sequences (Yaari et al., 2013). The second model (“RS5NF”) similarly assesses 5-mers but uses both synonymous and non-synonymous (replacement) mutations from non-productively (non-functional) rearranged sequences (Cui et al., 2016). Such models have been used to simulate B cell repertoire lineages by constructing a set of hypothetical sequences that have been mutated in a sequential manner as governed by, for example, the underlying S5F substitution scores (Krantsevich et al., 2021; Sheng et al., 2017). Although *k*-mer approaches are generally able to capture some key local intrinsic biases of SHM, such as hotspot targeting, there is evidence that shorter *k*-mers are insufficient to properly characterize differential SHM targeting. For example, a recent study extended a local sequence (5-mer) context model and improved accuracy by including parameters describing the position within the IGHV gene (Spisak et al., 2020). Another study compared the mutability of identical 5-mer (middle position +/-2nt) motifs at different positions within an IGHV gene (Zhou and Kleinstein, 2020), and found that the mutation frequency of these motif-allele pairs (MAPs) positively correlates with the overall mutability of a wider neighborhood of motifs, suggesting that an extended *k*-mer may better capture SHM.

Earlier studies have shown that using deep learning is effective in different genomic applications; for example, convolutional neural networks (CNNs) in extracting conserved sequence motifs among target sequences (Alipanahi et al., 2015; Kelley et al., 2016; Zhou and Troyanskaya, 2015). In this study, we adopted a deep learning approach using a 2-D CNN to analyze extended *k*-mer lengths to better understand the underlying SHM process. We demonstrate that our model, DeepSHM (Deep learning on SHM), can more accurately represent the SHM process by evaluating longer *k*-mers of up to 21 nts. Additionally, DeepSHM using 15-mers as inputs was able to recapitulate AID WR<u>C</u>/<u>G</u>YW hotspot motifs and identify an extended DWR<u>C</u>T motif. Neural network predictions are notoriously difficult to explain (the “black box” problem), but many new methods are available to interpret results (Koo and Ploenzke, 2020). We used one such method to identify a negative association between increased mutability at a site and its surrounding G content. On the other hand, lower mutation frequency was correlated with increased substitution rates of certain substitution types, particularly for G>T and C>A mutations. Furthermore, many highly conserved sites within G-rich sub-regions belonging to several IGHV3 genes display an extremely high bias towards creating G mutations, some of which may participate in the formation of G-quadruplex (G4) structures.

## Results

### Deep learning can more accurately represent SHM mutabilities and substitution biases

The objective of our analysis was to use supervised deep learning to build an accurate convolutional neural network (CNN) for SHM and, as much as possible, identify novel features contributing to mutability. We chose CNNs because we still expected mutation frequency to depend on recurring motifs that might occur at any position in the sequence (most obviously, AID hotspots), a task CNNs are well suited to. The workflow of our network consists of an input layer that processes a *k*-mer subsequence represented in its one-hot encoding format (i.e. a 4×*k* matrix of zeros and ones), followed by a convolution layer and two fully connected layers as the hidden layers, and finally the output layer of size 4×1 or 1×1, depending on the task that is being predicted (**Figure 1**, see Methods). Several hyperparameters, including dropout rate and learning rate, were fine-tuned with our model as well (**Supplementary Table 1**). We defined a model that would separate mutations on each strand (which are predominantly at C and A on the top strand and at G and T on the bottom strand) at the input level. To achieve this, we identified a simple solution using padding that assigns a row in each channel of the convolution layer output separately to each strand (**Figure 1**). CNNs are also often used together with attribution methods such as Integrated Gradients, to help with interpretation of the results.

**Figure 1.**
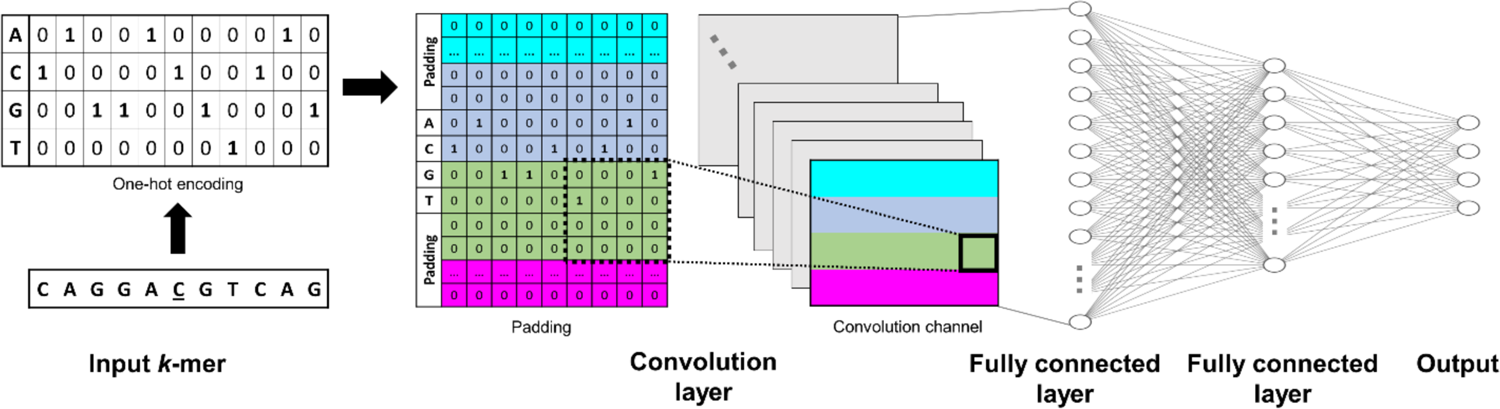
DeepSHM model architecture. Each model had an input layer, one convolution layer, two fully connected layers, and an output layer. The input layer was a 4×*k* dimensional one-hot encoded matrix (*k* is length of subsequence). The dimension of the output layer was dependent on the task: substitution (4×1), mutation frequency (1×1), or weighted substitution (4×1). For the convolutional layer, ‘same’ padding was used to allow the model to process top and bottom strand mutations separately. With ‘same’ padding, the output of each convolutional channel has the same shape as the input (4×*k*) with the following properties: the first and the fourth rows are populated with zeros only (there was no real input, only padding; cyan and magenta rows); the input used for the second (light blue) row contained two rows of padding and two data rows corresponding to A or C nucleotides only; and similarly, the input used for the third (green) row also contained two rows of padding and two rows of data corresponding to G or T nucleotides. Since AID and Polη target C and A sites respectively, this approach was taken with the expectation of helping the model distinguish top and bottom strands.

As a starting point, we trained two CNN models, which we collectively refer to as DeepSHM (Deep learning on SHM), to separately predict mutation frequency and substitution rates, calculated from previously published B cell repertoire data containing non-productively rearranged and clonally independent VDJ coding sequences (Tang et al., 2020), for varying *k*-mer lengths (see Methods). We trained both models independently using different combinations of *k*-mer lengths and hyperparameters as listed in **Supplementary Table 1**. We found that for predicting mutation frequency, 15-mers were moderately better than 9-mers (purple boxplots in **Figure 2A**, Mann-Whitney U test: P<2.2×10^-16^) and that further extending the motif length to 21 did not improve accuracy since both produced an overall maximum correlation (across hyperparameters) of 0.76 (**Figure 2A, Table 1**). Thus, using *k*-mers of length 15 or longer outperformed shorter lengths, specifically 5-mers and 9-mers (**Table 1**), suggesting that an extended DNA motif can better model the SHM process. However, using longer *k*-mers did not substantially improve the model that predicts SHM substitution bias alone, achieving an average correlation of 0.55 for 15-mers (green boxplots in **Figure 2B, Table 1**), but which is similar for different lengths. For the interpretability analysis below, we chose to use the best 15-mer models to keep the *k*-mer length consistent for comparisons across all models. In order to check if the performance of the models leading to the best results was consistent, we also trained 30 different iterations of each model, keeping the hyperparameters fixed but using different random seeds. We found the standard deviation across correlations was very small, at 0.002 for the mutation frequency model and 0.001 for the substitution rate model, showing the strong consistency of our results.

**Figure 2.**
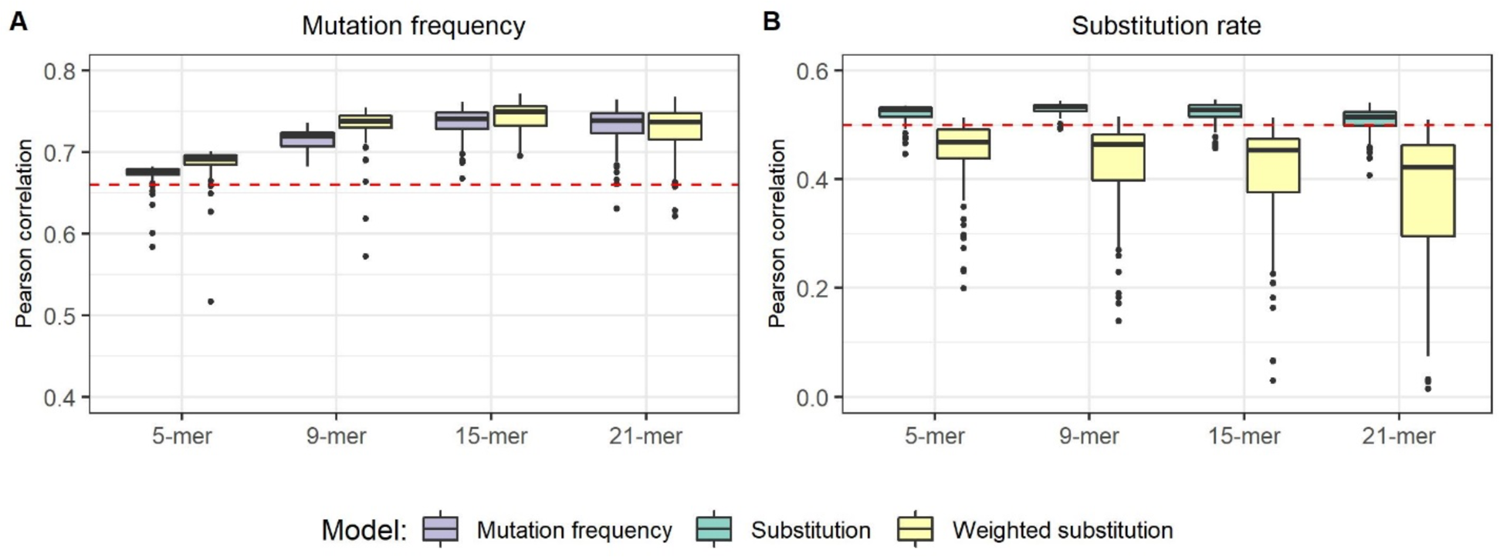
Performance of DeepSHM. Boxplots describing the distribution (across random hyperparameters) of Pearson correlations between DeepSHM predictions and empirical data (y-axis) are shown for different input *k*-mers (x-axis) for **(A)** mutation frequencies, and **(B)** substitution rates, for all three models (mutation frequency, substitution, and weighted substitution). Red dashed lines signify correlations of predicted S5F values, which uses 5-mers.

**Table 1.**
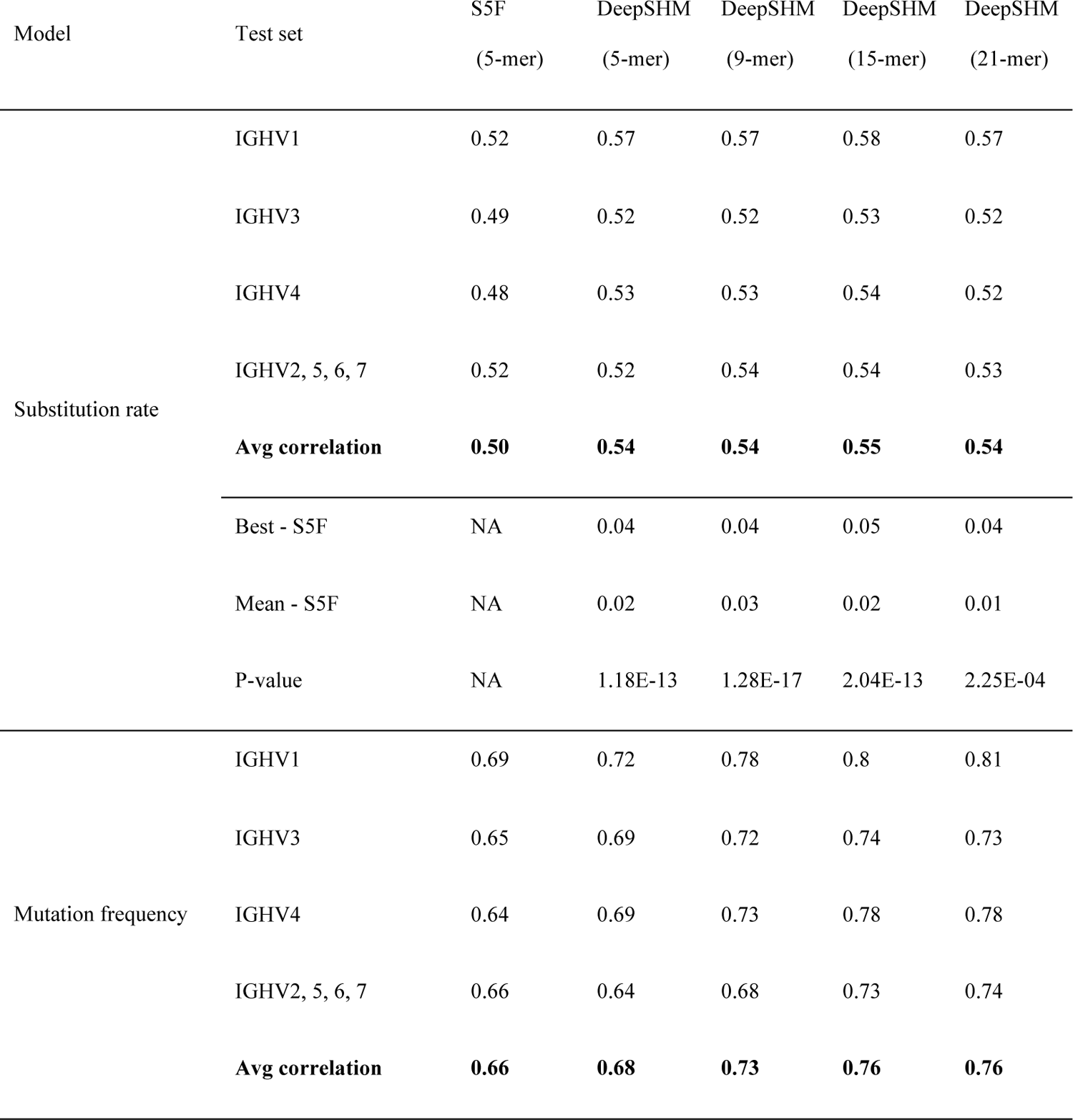
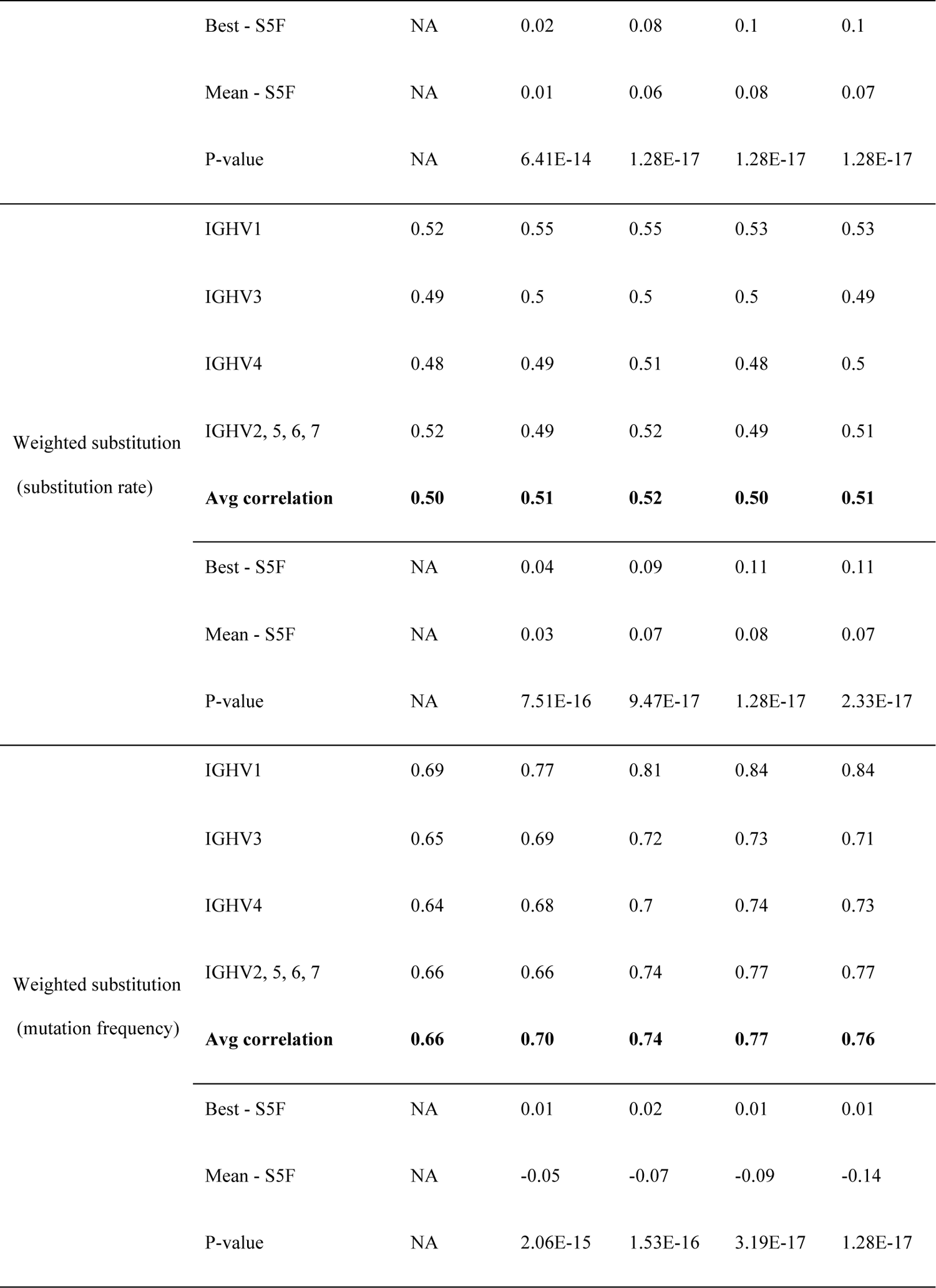
Cross-validation of various input *k*-mer sequences. The correlations of repeatedly trained models using different random seeds (but the same hyperparameters) for neural network training had small standard deviations, in all cases below 0.01. P-values are from a Wilcoxon signed-rank test comparing the training results for each model with the corresponding S5F model accuracy. P-values were corrected (Benjamini-Hochberg) for multiple comparisons.

We next sought to compare DeepSHM against the widely used S5F model that is based on 5-mer motifs (Yaari et al., 2013). To ensure a fair comparison, we generated an S5F targeting model using the same data set that was used to train DeepSHM, as well as the same cross-validation scheme (see Methods). Using the same test set splits as above, we found that there was an average correlation of 0.66 between the predicted S5F model mutabilities and empirical mutation frequency, and an average correlation of 0.50 for predicted S5F substitution scores and empirical substitution rates (red dashed lines in **Figure 2, Table 1**). The substitution model slightly (but statistically significantly) outperformed S5F for all *k*-mer values we analyzed. The mutation frequency model achieved a modest improvement over S5F using 5-mers as an input, and this difference became more evident for 9-, 15-, and 21-mers (**Figure 2A, Table 1**). We also similarly computed 30 iterations (using different random seeds) of the best 15-mer models for both mutation frequency and substitution models, and found these iterations to have significantly greater accuracy than S5F both individually and in aggregate (P<1.8×10^-6^ for each model, Wilcoxon signed-rank test). Overall, these results show that our deep learning approach successfully extracts meaningful information from the wider sequence context to improve predictions.

To identify associations between mutation frequency and specific substitutions, we further constructed a DeepSHM model to predict the “weighted substitution” of a *k*-mer, i.e., the product of the percentage of each observable substitution type (e.g G>N) and the mutation frequency of the *k*-mer (see Methods). Note that this weighted substitution metric is a vector representing the four ordered DNA bases, with a “0” placed at the position that matches the middle nucleotide of the *k*-mer. Since weighted substitution constitutes aspects of both the observable mutation frequency and substitution rate of the middle nucleotide of a given *k*-mer, we were able to evaluate DeepSHM on each metric separately. Although this model made poorer substitution rate predictions on average (varying hyperparameters) than S5F (**Table 1**), the best model performed similarly to S5F for substitution rates while, surprisingly, performing slightly better than any model in predicting mutation frequency. Cross-validation in this instance produced a range of average correlations between 0.50-0.52 for predicted substitution rates – a level similar to that of S5F (**Figure 2B, Table 1**). On the other hand, DeepSHM of weighted substitution values was marginally better at predicting mutation frequency for 15-mers (correlation: 0.77) than the previous standalone model that was tasked to learn mutation frequency only as well as being better than S5F. (**Figure 2A, Table 1**). Since the weighted substitution model was able to perform at a level slightly better to the standalone mutation frequency model for longer *k*-mers and substantially better for shorter (5-mer, 9-mer), this suggested a possible association between the projected substitution bias of a site and overall mutability and furthermore, that interpretability methods might uncover these (see below).

### Interpretation of the DeepSHM network reveals extended hotspot motifs

A complication often associated with deep neural networks is model interpretability (the “black box” problem). One way we interrogated the predictions made by DeepSHM, and what it has learned about the SHM process, was to analyze the output of the penultimate layer of each 15-mer based model. In particular, analyzing the output, or “encodings”, of this layer can be viewed as an alternative, and more informative, way of representing the input 15-mer. To visualize the multi-dimensional encodings of the individual 15-mers, we used t-SNE, a dimensionality reduction technique, to project each onto a 2-dimensional embedding (see Methods). At this point in order to make full use of the data, we merged all of the 15-mer data into one training set, and then trained three new individual models (one for each output type) using the hyperparameters which previously led to the best cross-validation results. The analyses we present below are derived from the DeepSHM models that were trained using this merged data set.

We began by identifying features learned by DeepSHM that predicted weighted substitutions. Since weighted substitution is a measure of both mutation frequency and substitution bias, the embedding should capture both metrics simultaneously. Each point in the resulting t-SNE embedding in **Figure 3A** represents a single 15-mer and is colored according to its corresponding mutation frequency. We identified several clusters of 15-mers that are mostly grouped by similar mutation rates, including those expressing high mutability. Clusters with mid to high mutation frequencies are similarly within close proximity but displayed no obvious groupings other than being mostly located towards the center. When we considered the middle nucleotide of each 15-mer, we observed that these clusters also shared the same middle nucleotide (**Figure 3B**), suggesting that the network identified as a key feature the “0” value in the weighted substitution output vector that is associated with the middle nucleotide.

**Figure 3.**
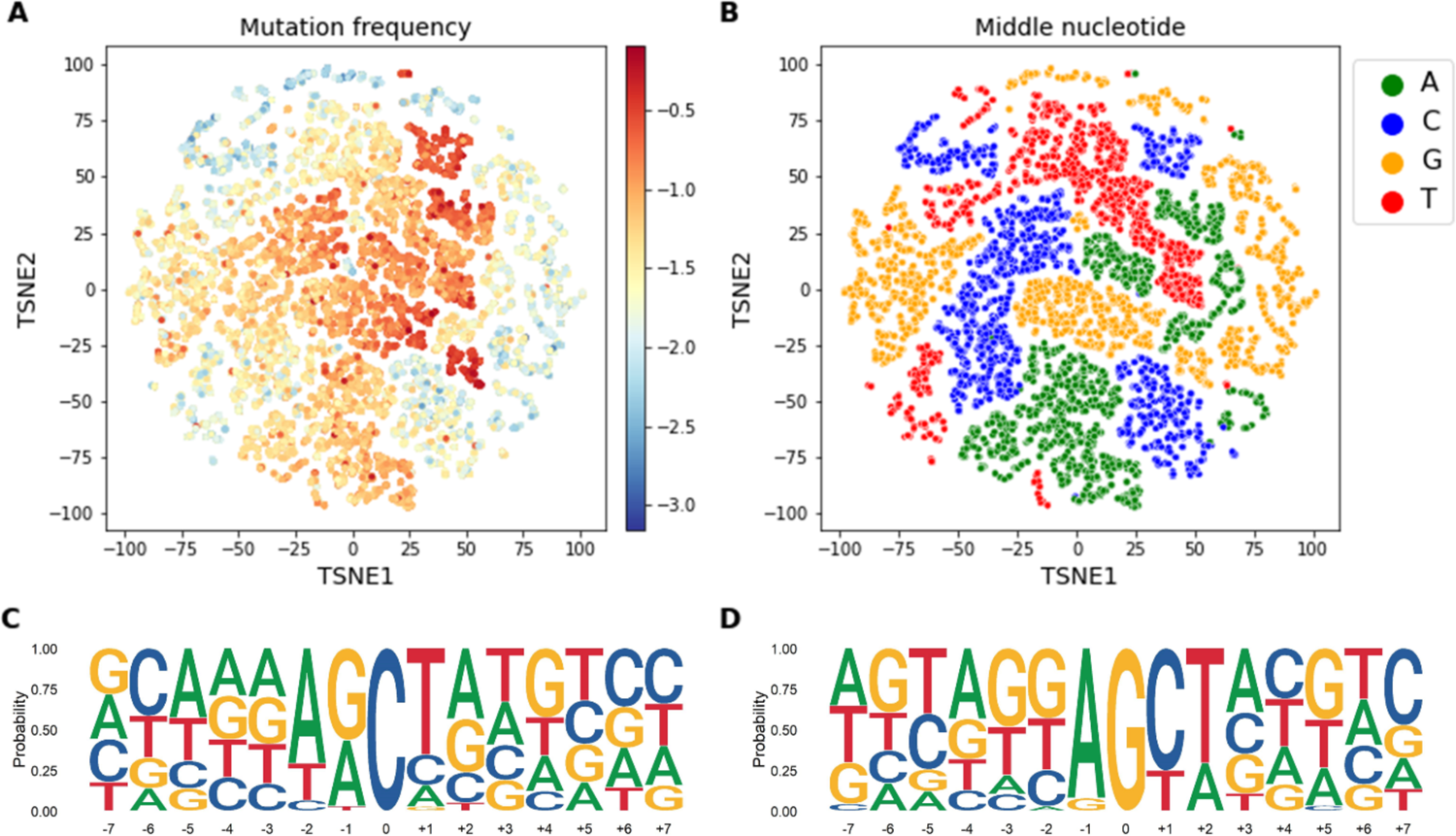
Neural network encodings analysis: weighted substitution model. Each point in the t-SNE embedding represents a single 15-mer processed through the truncated model (to extract the output of the penultimate layer) originally trained to learn the associated weighted substitutions (see Methods) and is colored according to its corresponding **(A)** mutation frequency (log_10_), and **(B)** middle nucleotide. Consensus sequences derived from the highest mutated cluster identified using k-means clustering on the embedding of 15-mers containing either **(C)** a middle C nucleotide or **(D)** a G nucleotide (clusters 10 and 16 in Supplementary Table 2).

Next, we applied k-means clustering on the embedding as a way to isolate cluster boundaries (**Supplementary Figure 1**, see Methods). We subsequently created a sequence logo plots representing each cluster shown in **Supplementary Table 2**. As expected, clusters with the highest mutation frequencies had inner subsequences containing AID (C cluster 10, G cluster 16) and Polη (A clusters 1 and 2, T cluster 20) hotspots. For AID, these are WR<u>C</u> (**Figure 3C**) and <u>G</u>YW motifs (**Figure 3D**). Within the two most highly targeted AID hotspot clusters there is a substantial presence of both WG<u>C</u>/<u>G</u>CW and WA<u>C</u>/<u>G</u>TW contexts, rather than only the well-known W<u>GC</u>W overlapping hotspot motif (Tang et al., 2020; Wei et al., 2015). Furthermore, even when we include WA<u>C</u>/<u>G</u>TW, there is a preference for a T base at the 3’ end of the WR<u>C</u> hotspot, and conversely, an even stronger bias for an A base at the 5’ end of the <u>G</u>YW hotspot (**Figure 3C, D**). This motif is consistent with a genome-wide study of AID mutations in mice that reported observing high mutability at AA<u>C</u>T and AG<u>C</u>T motifs in both strands (Álvarez-Prado Á et al., 2018). When we assessed the mutability of all possible WRCN (N=A/C/G/T) motifs separately, we observed WRCT to be the most highly mutated in each case (**Supplementary Figure 2**). Previous studies identified WR<u>C</u>Y/R<u>G</u>YW (Y=C/T, R=A/G) and later WR<u>C</u>H/D<u>G</u>YW (H=A/C//T, D=A/G/T) to be a better predictor of mutability at C:G bases (Rogozin and Diaz, 2004; Rogozin and Kolchanov, 1992). However, we discovered some inconsistencies with these definitions, as AG<u>C</u>C was found to be the least mutated of the AG<u>C</u>N motifs and WR<u>C</u>G was not always the least mutated, on both strands. Overall, these early hotspot definitions may have been too broad, and WR<u>C</u>T/A<u>G</u>YW is a more consistent predictor of AID targeting. Lastly, we also noted that among the least mutated *k*-mer clusters, many were G-rich in their surrounding context (for example, C clusters 8 and 9, A cluster 4, T cluster 21), and particularly for G (G clusters 12 and 14) (**Supplementary Table 2**).

As a complementary way to find sequence motifs associated with mutability, we used TF-MoDISco (Transcription Factor Motif DIScovery), a program for identifying recurring motif patterns in genomic data (see Methods) (Shrikumar et al., 2018). We applied TF-MoDISco to the standalone model that predicts only mutation frequency because we reasoned that sequence features related to mutability would be more easily identifiable since the model is only required to learn a single task. TF-MoDISco uses importance scores, which can be derived from many machine learning methods, to produce a set of unique motifs learned by the model (see Methods).

We began by analyzing the importance scores derived from Integrated Gradients (Sundararajan et al., 2017) of 15-mers whose middle nucleotide conformed a WR<u>C</u>/<u>G</u>YW AID hotspot motif. As expected, the positively contributing sites in the set of ensuing motifs aligned with the hotspot motifs (**Figure 4A, B**). In addition, TF-MoDISco again revealed a preference for having a T base at the +1 position of the WR<u>C</u> (WR<u>C</u>T, **Figure 4A**) and an A base at the −1 position of the <u>G</u>YW (A<u>G</u>YW, **Figure 4B**).

**Figure 4.**
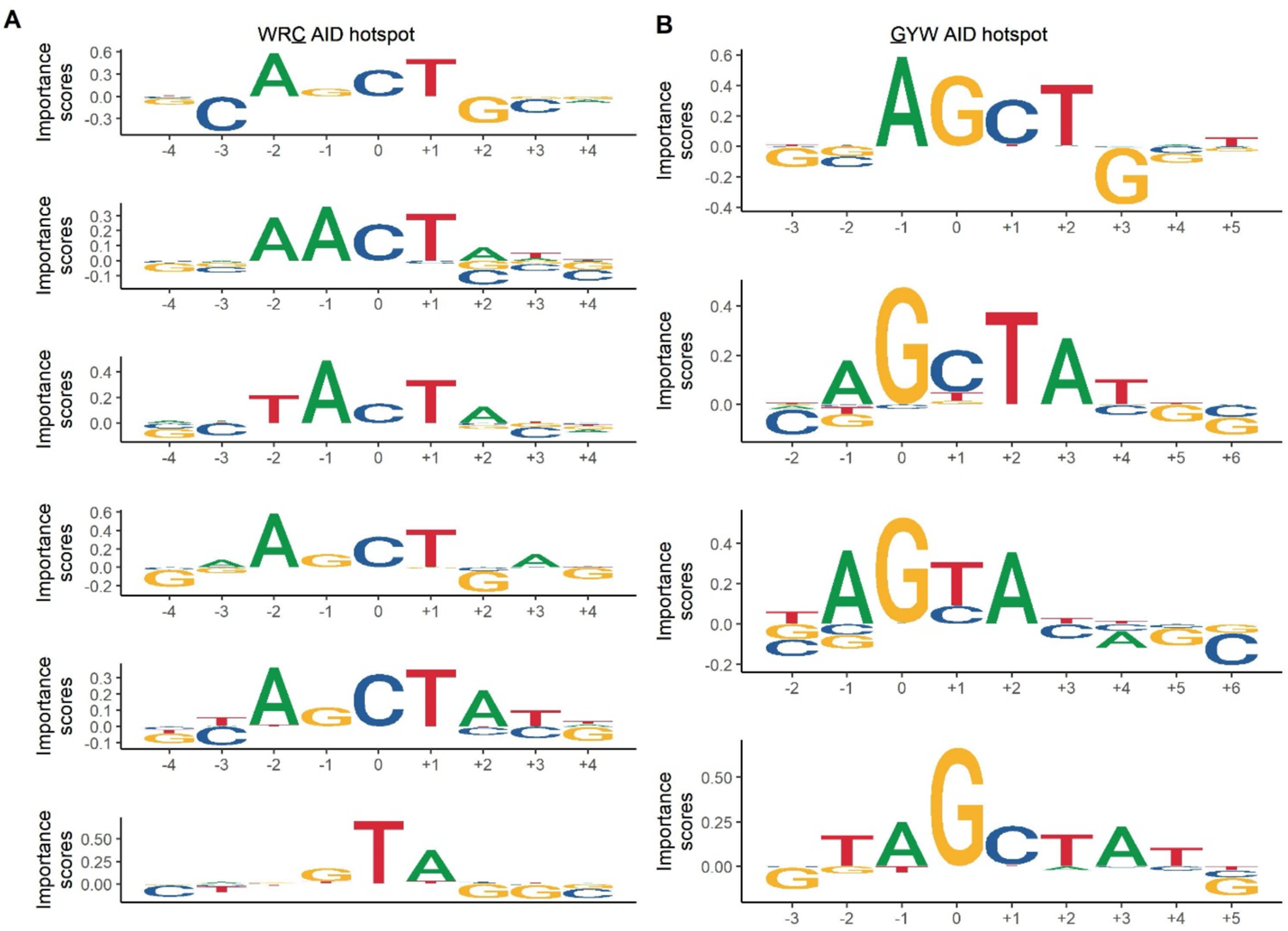
Recurrent motifs identified by TF-MoDISco. TF-MoDISco results using the Integrated Gradients as base-level importance scores of 15-mers whose middle nucleotide conformed to a **(A)** WR<u>C</u> or **(B)** <u>G</u>YW AID hotspot motif.

In addition to WR<u>C</u>T/A<u>G</u>YW being a well-represented motif identified by TF-MoDISco, as measured by having positive contributions to mutability (above horizontal axis on **Figure 4A**), we also noticed many neighboring C and G bases contained negative contributions (below horizontal axis on **Figure 4A**), most evidently at the C located at the −3 position of the WR<u>C</u> hotspot, and the G located at the +3 position of the <u>G</u>YW hotspot (**Figure 4B**). Here, the negative contribution at the −3 position signifies that having a C at that position reduces mutational targeting to the middle C. By the same token, a mutation that changes the −3 position from C will increase the likelihood of the middle C subsequently being targeted. This observation supports our recently published study where we observed a strong positive “mutual association” – a correlation metric describing the impact of mutating one site and its effect at another site – between CC (or GG) pairs distanced by two nucleotides (Krantsevich et al., 2021). In that study we were able to explain most of such correlations in terms of overlapping AID and/or Polη hotspots, with the CNNC/GNNG motif being one the exceptions which we suggested might be explained by AID processivity (Pham et al., 2003; Storb et al., 2009). However, the TF-MoDISco analysis suggests a different explanation in which the absence of a C in the −3 position might be a part of an extended AID hotspot, defining <u>C</u>WR<u>C</u> as being similar to a sequential overlap motif, which we previously defined (Krantsevich et al., 2021) as a motif in which an initial mutation creates a new hotspot that previously did not exist. Here, although the WR<u>C</u> hotspot did previously exist, a mutation in the first C would create a DWR<u>C</u> (D=A/G/T) motif, potentially with higher mutability.

We next sought to determine whether adding the 5’ D or 3’ T context of the canonical WR<u>C</u> hotspot is more influential in terms of increasing its susceptibility to AID mutagenesis. To address this, we increased the hotspot specificity step by step, starting from CWR<u>C</u>V (V=A/C/G) and assessed the impact a single change in the motif at either the first C or V site, causing a DWR<u>C</u>V or CWR<u>C</u>T intermediate hotspot to form respectively, has on mutability (**Figure 5A**). We found that both DWR<u>C</u>V and CWR<u>C</u>T intermediate hotspots were shown to mutate significantly more than CWR<u>C</u>V (**Figure 5B**). We also discovered that the mutability of the DWR<u>C</u>T hotspot, which contains the extended hotspot in both 5’ and 3’ directions, was significantly higher than both intermediate hotspots (**Figure 5B**). Performing a pairwise comparison between the mutation frequency of all 16 individual (D/C)WR<u>C</u>(T/V) contexts further confirmed that those containing both a 5’ D and 3’ T were significantly more mutated than the remaining hotspot motifs, with DAG<u>C</u>T being the most mutated (**Figure 5C, Supplementary Table 3**). Additionally, the next three successively mutated hotspots followed a CWR<u>C</u>T context, overall suggesting the 3’ T to be more impactful to AID recognition than the 5’ D, but the addition of both substantially increases targeting in human V regions.

**Figure 5.**
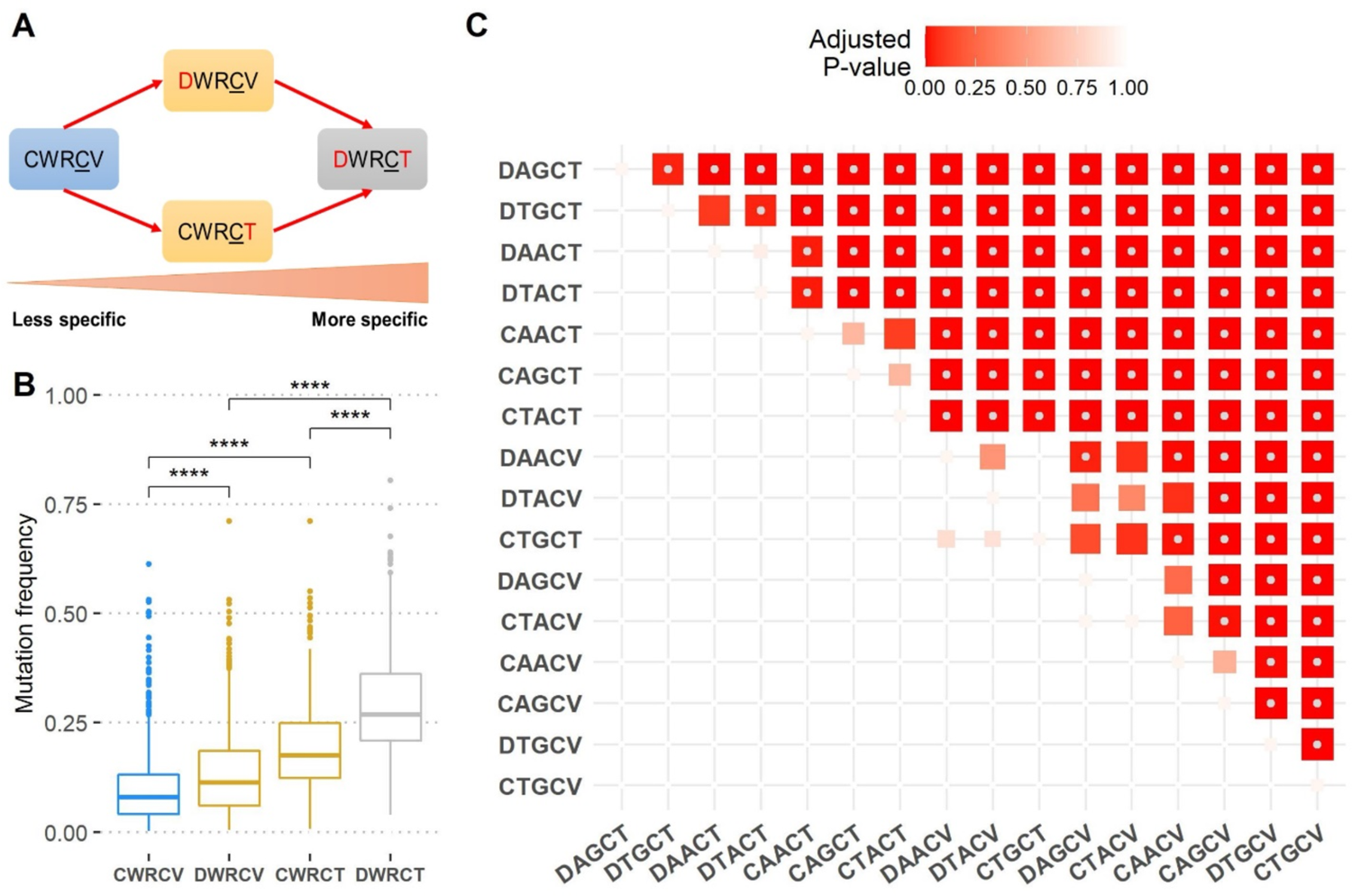
Mutability of extended AID hotspots. **(A)** Schematic showing an increase of AID hotspot specificity (left to right). **(B)** Boxplots displaying the mutability of different (C/D)WRC(T/V) hotspot contexts, where D=A/G/T, V=A/C/G. Asterisks indicate significance (p ≤ 0.0001) of a one-sided Mann-Whitney U test comparing the greater mutation frequency of the boxplot on the right against the one on the left. **(C)** Pairwise comparison of mutability for all 16 (C/D)WRC(T/V) hotspot contexts. Boxes represent the p-value - adjusted for multiple comparisons (Benjamini-Hochberg correction) - of a one-sided Mann-Whitney U test comparing the greater mutation frequency of the hotspot indicated by the row to the left, against the hotspot shown in the column below. Rows and columns are ordered by mean mutation frequency (high to low). The color and size of each box is scaled according to the adjusted p-value. Gray dots inside boxes indicate p-values ≤ 0.05.

In addition, another secondary motif that unexpectedly emerged from the TF-MoDISco analysis of WR<u>C</u>/<u>G</u>YW 15-mers did not contain a positively contributing C nucleotide; rather it conformed to a <u>TA</u> Polη hotspot (**Figure 4A**, bottom). Having a <u>TA</u> hotspot appear while specifically analyzing only 15-mers containing WR<u>C</u> hotspots reveals the importance of attracting Polη to these areas. This finding is consistent with our previous analysis highlighting the importance of co-localization of A<u>GC</u>T overlapping AID hotspots and Polη hotspots within the CDRs (Tang et al., 2020; Wei et al., 2015).

The <u>TA</u> motif also emerged when we applied TF-MoDISco to all 15-mers conforming to either a W<u>A</u> (**Supplementary Figure 3A**) or <u>T</u>W Polη hotspot (**Supplementary Figure 3B**). In addition to our model identifying both the <u>TA</u> and A<u>A</u> hotspot motifs as important, it also identified a T<u>A</u>T/A<u>T</u>A motif as a special case for both strands. Further analysis showed that W<u>A</u>T/A<u>T</u>W hotspots mutate significantly more than their W<u>A</u>V/B<u>T</u>W counterparts (**Figure 6**). Thus, while T<u>A</u> hotspots consistently have higher mutability than A<u>A</u>, the presence of a 3’ T individually increases the mutability of each of these Polη hotspots.

**Figure 6.**
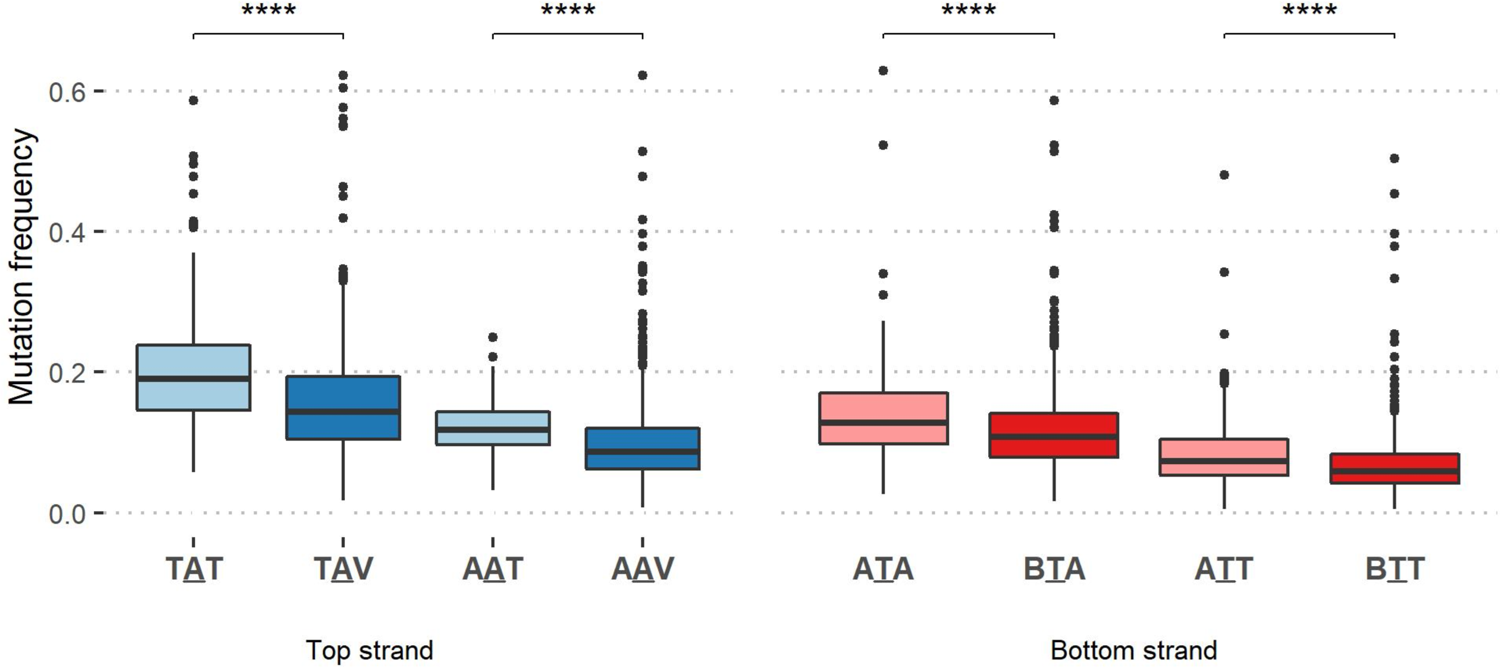
Mutability of extended Polη hotspot motifs. Boxplots comparing the mutation frequency of various top strand W<u>A</u>T against W<u>A</u>V (blue), and bottom strand A<u>T</u>W against B<u>T</u>W (red) motifs. Asterisks indicate significance (p ≤ 0.0001) of a one-sided Mann-Whitney U test comparing the greater mutation frequency of the boxplot on the left against the one on the right.

### Highly targeted sites display a lower surrounding GC content

We next applied the same t-SNE methodology to the DeepSHM model that predicted only mutation frequency. We found that the organization of the subsequent embedding followed a direction of descending mutation frequency, with the highest mutating 15-mers located at the mid- to upper-right portion of the plot (**Figure 7A**). A cluster of low-mutating 15-mers was also isolated in the upper-left (**Figure 7A**), which was enriched with ∼76% of FW1 15-mers (**Figure 7B**). Additionally, we examined the possible influence of the local surrounding sequence by calculating the individual base content of the four DNA bases in each 15-mer. However, the inner 5-mer, which contains the dominant context, was excluded when computing all base counts. When we colored the t-SNE embedding according to the GC content of each 15-mer, we observed that GC content increases along the same direction as decreasing mutability seen previously (**Figure 7C**). Quantifying this observation more formally, we indeed found a significant negative correlation between the GC content and the mutation frequency of the 15-mers (R=-0.31, P<2.2×10^-16^; **Figure 7D**). On the other hand, when we considered each individual base count independently, we observed that the count of G nucleotides specifically shows a stronger negative correlation (R=-0.19) than the C nucleotide count (R=-0.084) alone (**Supplementary Figure 4A**), although both correlations are highly significant (P<2.2×10^-16^). This result is consistent with the cluster analysis above (**Supplementary Table 2**) where we observed several clusters with G-rich *k*-mers and low mutation frequencies. If we further separate the mutation frequencies into categories defined by the middle nucleotide, we find that G content has a consistent negative correlation regardless (column G of **Supplementary Figure 4B**). More generally, A and T richness (columns A and T of **Supplementary Figure 4B**) shows a consistent positive or sometimes non-significant correlation, whereas C and G richness shows a consistent negative (or non-significant) correlation. In summary, it appears that low-mutating sites generally have a high local GC (and particularly G) content, and conversely, that highly targeted sites display an elevated local AT (particularly A) content.

**Figure 7.**
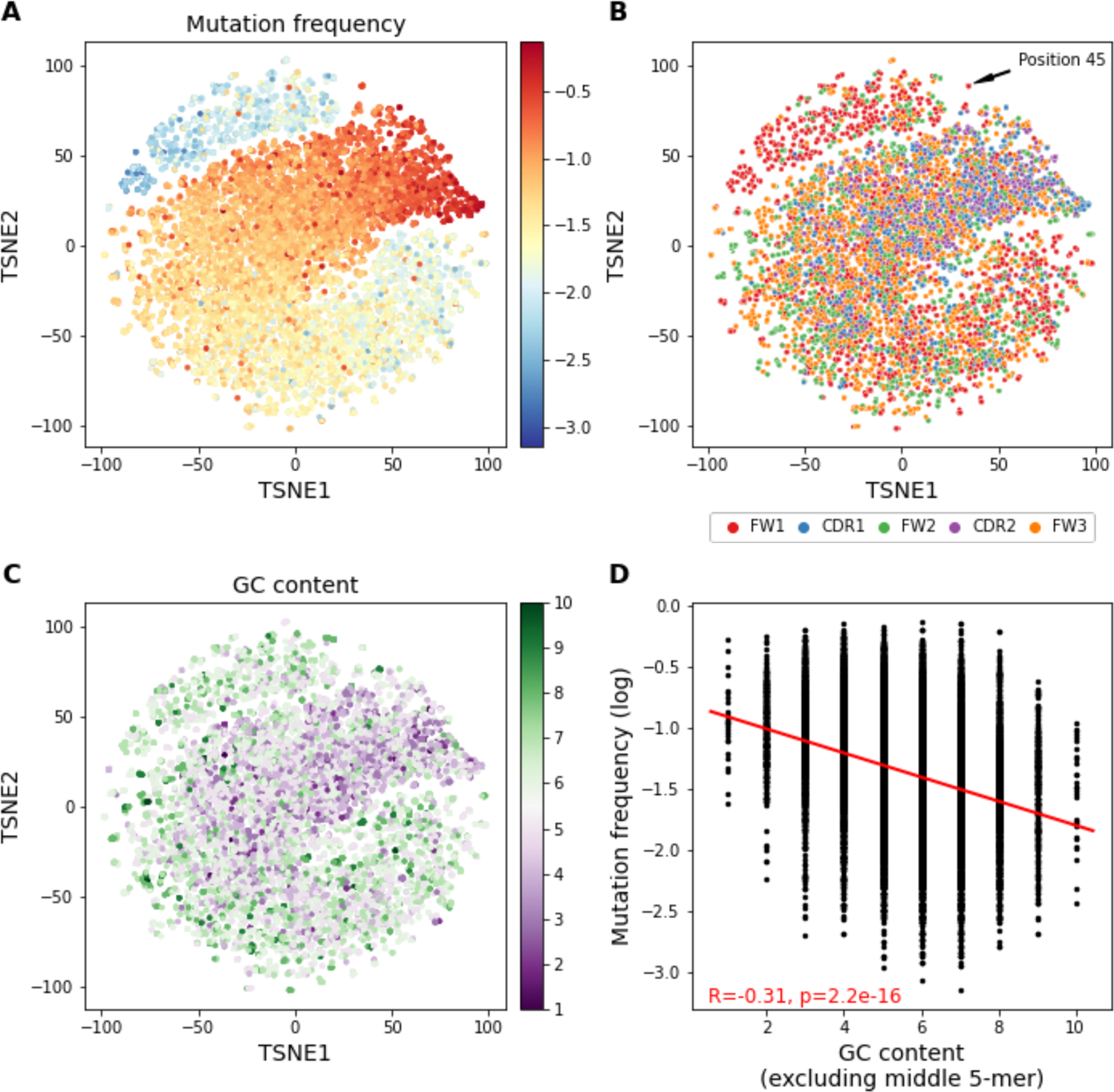
Neural network encodings analysis: mutation frequency model. Each point in the t-SNE embedding represents a single 15-mer processed through the truncated model (to extract the output of the penultimate layer) trained on mutation frequencies (see Methods) and is colored according to its corresponding **(A)** mutation frequency (log_10_), and **(B)** by Ig V sub-region location as defined by IMGT. **(C)** The t-SNE embedding is colored according to the GC content of each 15-mer. The calculated GC content excludes the middle 5-mer context of the 15-mer to remove any confounding AID hotspot or coldspot bias. **(D)** Computed Pearson correlation between mutation frequency and GC content, again excluding the middle 5-mer.

### Conserved FW1 sites surrounded by clusters of AID coldspots in IGHV3 genes display a high T>G transversion bias

We now analyzed the standalone model predicting only substitution rates to gain possible insight into additional substitution biases exhibited by AID or downstream error-prone DNA damage response pathways, for example, as a result of REV1 or Polη intervention during non-canonical base-excision repair (BER) and non-canonical mismatch repair (MMR), respectively. The resulting t-SNE embedding from this model identified four main clusters, as well as two much smaller satellite clusters, with each cluster containing 15-mers that share a common middle nucleotide (**Figure 8A**). A distinction between 15-mers with high and low mutation frequencies could also be observed based on their location on opposite ends of the cluster, especially for clusters containing either a C or G middle nucleotide, with high-mutating 15-mers typically located on the side closest to the center (**Figure 8B**). Since the model was tasked with learning the distributed substitution rates of each 15-mer, we next sought to evaluate the embedding by the rate of each individual substitution type (e.g. C>T). In certain clusters, a similar gradient of high to low substitution rates could also be seen as we observed for mutation frequency (**Figure 8C-F**). For instance, we noticed the rate of G>T substitutions increasing from the side nearest to the origin towards the outer boundaries of the cluster (top-right cluster in **Figure 8F**), which was associated with a shift towards decreasing mutation frequencies in the same cluster while proceeding in the same direction (**Figure 8B**). To evaluate this trend more closely, we analyzed three human IGHV genes from different families for which we had the most data (IGHV1-18, IGHV3-23, IGHV4-34), so as to include sites with low mutation frequencies at high coverage, and calculated the correlation between mutation frequency and rate of substitution for each substitution type. As an example, for IGHV3-23 we found the most significant negative correlations to be at C>A mutations (R=-0.33, p=0.0058), and the reverse, G>T (R=-0.24, p=0.022; **Supplementary Figure 5**). Alternatively, we observed a significant positive correlation between mutation frequency and C>T transition mutations (R=0.29, p=0.018; **Supplementary Figure 5**). Similar patterns were also observed for IGHV1-18*01 and IGHV4-34*01 (**Supplementary Figure 5**). These results are consistent with replication bypass (predominantly causing C>T) being favored over BER at sites with high mutation frequency.

**Figure 8.**
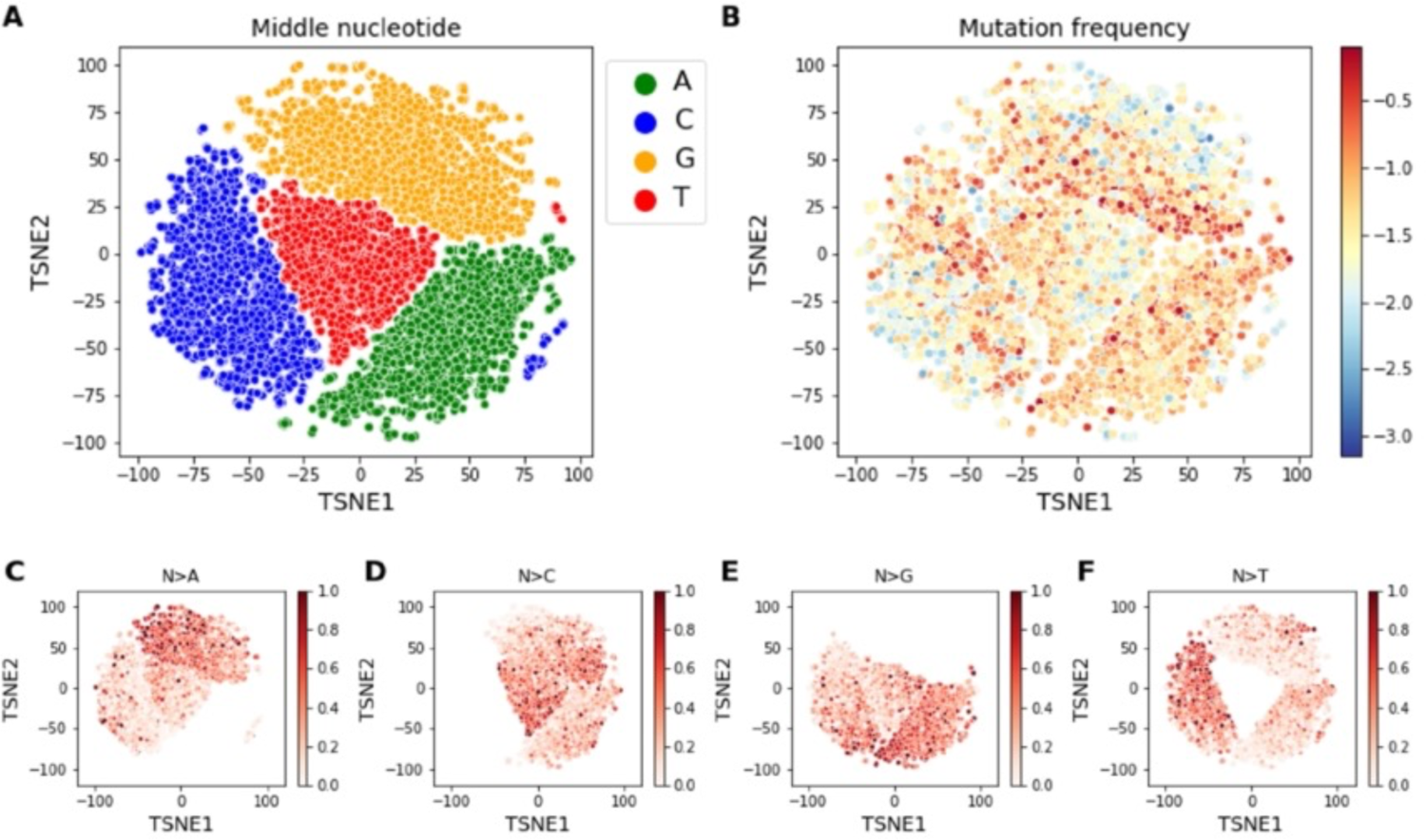
Neural network encodings analysis: substitution model. Each point in the t-SNE embedding represents a single 15-mer processed through the truncated model (to extract the output of the penultimate layer) trained to learn the associated substitution rates (see Methods) and is colored according to its corresponding **(A)** middle nucleotide, and **(B)** mutation frequency (log_10_). **(C-F)** The t-SNE embedding is colored by the rate of substitution for the middle nucleotide of every 15-mer to mutate to A (N>A); to C (N>C); to G (N>G); and to T (N>T), respectively.

In the t-SNE analysis of the substitution model, we also discovered two small clusters of 15-mers containing a C and T as their middle nucleotide (**Figure 8A**) that did not group with their respective larger clusters, suggesting that these particular sites might have distinct substitution patterns. Generating the consensus sequence of the outlier T cluster revealed a partially conserved AGYC<u>T</u>GGGGG sequence (**Figure 9A**). When we examined these subsequences more closely, we discovered that they were located only in IGHV3 family genes at either position 21 or position 45 according to the IMGT unique numbering system (Lefranc, 2001) (**Supplementary Figure 6**). The motif was also surprisingly common. At position 21 it appeared in 37 different alleles (across 19 genes) and was fully conserved in all alleles. Coincidentally, the motif also appeared in 37 different alleles (across 18 genes) at position 45, although it differed slightly at the +3 and +4 positions (**Figure 9A**). These two sets of alleles only partially overlap, such that 15 alleles had the motif at both positions 21 and 45. Thus, this specific motif in FW1 of the IGHV3 family genes appears to be highly conserved evolutionarily, suggesting a possible functional role. The rates of substitution at these sites were also found to be highly biased towards creating T>G mutations, with an average T>G rate of about 0.66 at position 21, and an even greater rate of 0.89 at position 45 (background rate: 0.28 ± 0.23) (**Figure 9B, Table 2**). A previous study using Sanger sequencing data that was limited to IGHV3-23 and the pseudogene IGHV3-h had noted similarly high T>G substitution rates at positions 21 (for IGHV3-h) and 45 (for IGHV3-23) (Ohm-Laursen and Barington, 2007). Although the T subjected to mutation at both positions did not conform to a bottom strand <u>T</u>W Polη hotspot, these genes at position 45 displayed a relatively high average mutation frequency of 0.17 ± 0.08 (**Table 2**), which is somewhat unusual given that mutations are generally more biased towards the CDRs than FW regions (Cohen et al., 2011; Shapiro et al., 2002), and that we reported above that many sites within FW1 tended to display low mutability (**Figure 7A, B**).

**Figure 9.**
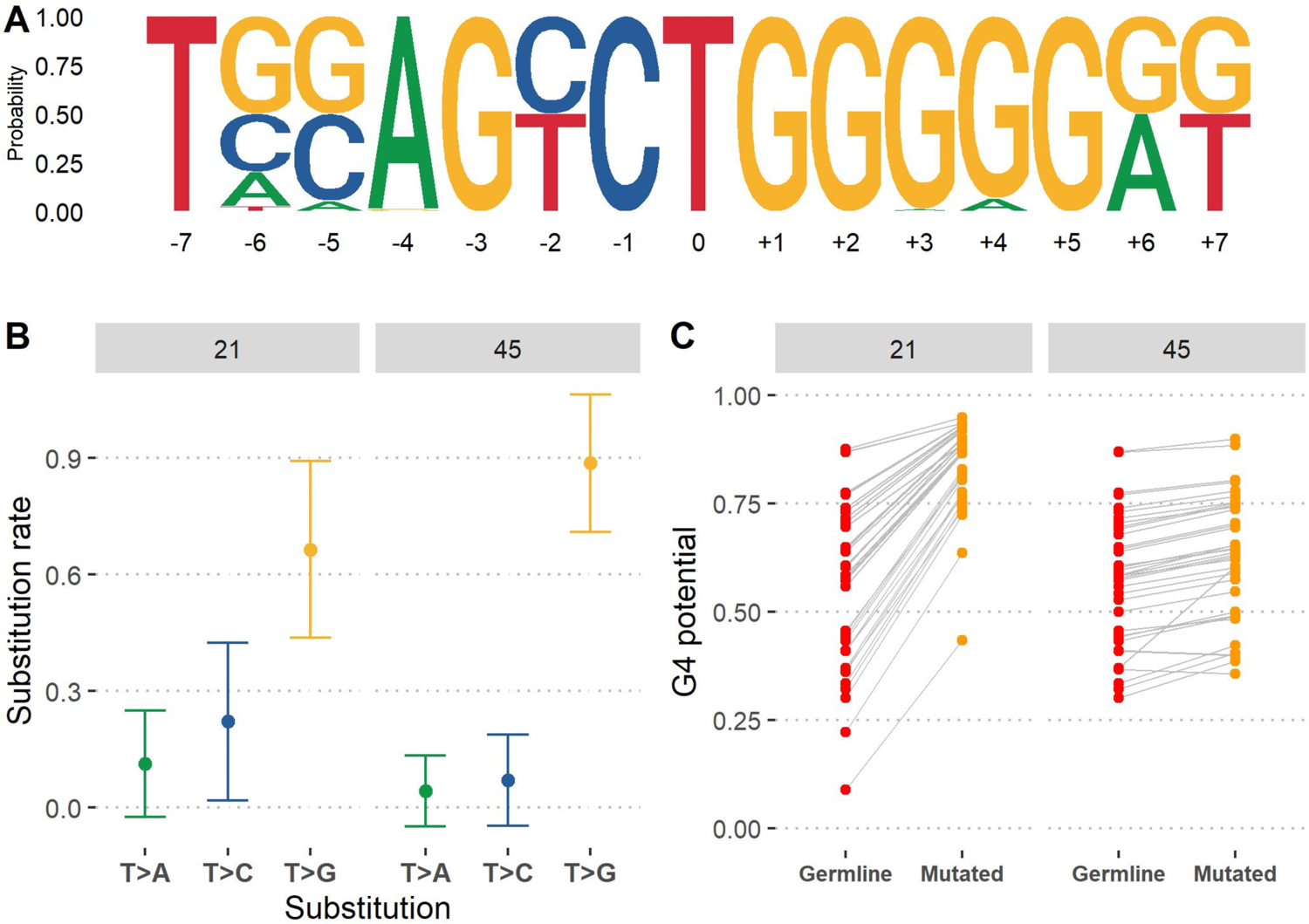
Evaluation of the T outlier cluster in the DeepSHM substitution model. **(A)** Sequence logo representation of the 15-mers appearing in the T outlier cluster in **Figure 8A** (right-hand side, red dots). **(B)** Substitution rates of T>A, T>C, and T>G for 15-mers corresponding to 37 IGHV3 alleles separately at IMGT positions 21 and 45. Bars represent ±1 standard deviation. **(C)** G-quadruplex (G4) formation potential for the same IGHV3 alleles in **(B)**. G4 potentials (y-axis) are computed using the germline IGHV sequence (“Germline”) and the mutated sequence (“Mutated”) containing a single simulated T>G mutation at either IMGT positions 21 or 45.

**Table 2.** Summary statistics on outlier C and T clusters from **Figure 8A**.

| IMGT position | 15-mer middle nucleotide | n | Avg. substitution rate to G | Avg. mutation frequency | Avg. germline G4 potential | Avg. mutated G4 potential | Avg. difference in G4 potential (mutated - germline) |
| --- | --- | --- | --- | --- | --- | --- | --- |
| 1 | C | 1 | 0.67 | 0.02 | 0.01 | 0.01 | 0.00 |
| 20 | C | 31 | $0.50 \pm 0.32$ | $0.01 \pm 0.01$ | $0.54 \pm 0.20$ | $0.64 \pm 0.19$ | $0.09 \pm 0.02$ |
| 21 | T | 37 | $0.66 \pm 0.23$ | $0.05 \pm 0.03$ | $0.54 \pm 0.19$ | $0.84 \pm 0.10$ | $0.29 \pm 0.10$ |
| 34 | C | 19 | $0.60 \pm 0.37$ | $0.03 \pm 0.05$ | $0.07 \pm 0.08$ | $0.07 \pm 0.08$ | $0.01 \pm 0.00$ |
| 44 | C | 38 | $0.77 \pm 0.24$ | $0.02 \pm 0.01$ | $0.56 \pm 0.17$ | $0.64 \pm 0.18$ | $0.07 \pm 0.04$ |
| 45 | T | 37 | $0.89 \pm 0.18$ | $0.17 \pm 0.08$ | $0.58 \pm 0.15$ | $0.63 \pm 0.14$ | $0.05 \pm 0.04$ |
| 48 | A | 1 | 0.79 | 0.11 | 0.37 | 0.56 | 0.19 |
| 49 | A | 5 | $0.79 \pm 0.12$ | $0.07 \pm 0.03$ | $0.37 \pm 0.07$ | $0.53 \pm 0.05$ | $0.16 \pm 0.02$ |
| 61 | C | 45 | $0.67 \pm 0.14$ | $0.04 \pm 0.02$ | $0.54 \pm 0.19$ | $0.54 \pm 0.19$ | $0.01 \pm 0.01$ |
| 101 | C | 1 | 0.52 | 0.11 | 0.01 | 0.01 | 0.00 |
| 167 | C | 1 | 0.46 | 0.52 | 0.37 | 0.41 | 0.04 |
| 173 | C | 3 | $0.64 \pm 0.29$ | $0.09 \pm 0.09$ | $0.04 \pm 0.02$ | $0.04 \pm 0.02$ | $0.00 \pm 0.00$ |
| 180 | C | 1 | 0.20 | 0.03 | 0.01 | 0.01 | 0.00 |
| 214 | C | 4 | $0.51 \pm 0.26$ | $0.13 \pm 0.06$ | $0.08 \pm 0.06$ | $0.10 \pm 0.07$ | $0.01 \pm 0.01$ |
| 249 | C | 15 | $0.59 \pm 0.24$ | $0.06 \pm 0.02$ | $0.08 \pm 0.09$ | $0.08 \pm 0.09$ | $0.00 \pm 0.01$ |
| 268 | C | 44 | $0.55 \pm 0.19$ | $0.06 \pm 0.03$ | $0.55 \pm 0.18$ | $0.53 \pm 0.18$ | $-0.01 \pm 0.01$ |

While examining the C outlier cluster (**Figure 8A**), we found the consensus sequence to be more diverse compared to the outlier with a middle T (**Supplementary Figure 7A**). The sequence variation seen here was partly due to the fact that the 15-mers that constituted this cluster belonged to many other IGHV families besides IGHV3 and across different sub-regions of the IgV (**Supplementary Figure 6**). On the other hand, we noticed some overlap between both outlier clusters since, in some cases, the C corresponded to positions 20 and 44 that preceded the middle T of the other outlier cluster (**Supplementary Figure 7A, Table 2**). We further found these sites to have a similar elevated C>G substitution rate (mean rate of 0.62 compared to background mean of 0.33, P<2.2×10^-16^) (**Supplementary Figure 7B, Table 2**), suggesting the model distinguished sites with a general preference to create G mutations.

Given that the sites with strikingly high T>G and C>G substitution rates we identified here are in adjoining G-rich sub-regions (**Figure 9A, Supplementary Figure 7A**), we evaluated the possible influence these mutations might have on the formation of G-quadruplex (G4) structures. In a recent study, we assessed the potential for DNA G4 structures to form in the IgV region, using a pre-trained deep learning model that computes the G4 potential of a linear DNA sequence (Tang and MacCarthy, 2021). There we found that the IGHV3 family had the highest propensity to form stable G4s in the top strand. We now sought to assess the overall mutational effect on G4 assembly of the IGHV3 sites that are biased towards G. Following the methodology of our previous study, we calculated the difference between the predicted G4 potential of the germline with that of the sequence with a single mutation at either position 21 or 45. Here, we found that a T>G mutation at position 21 elevated average G4 potentials to a very high value of 0.84 ± 0.10 compared to a germline value (already relatively high) of 0.54 ± 0.19, whereas the same mutation occurring at position 45 displayed a far smaller average increase of 0.05 ± 0.04 (**Figure 9C, Table 2**). As for the remaining cases, there seemed to be little effect of C>G mutations on G4 potential (**Table 2**).

Interestingly, we made another observation regarding the instances where an A nucleotide disrupts the run of G nucleotides at the +3 or +4 positions (**Figure 9A**) which was that these sites also displayed high A>G substitution rates (0.79 ± 0.11; **Table 2,** positions 48 and 49). This hypothetical mutation also caused a moderate, though substantial, increase in G4 potential (0.17 ± 0.02, **Table 2**). These findings reveal that particular recurring mutations in this sub-region may promote G4 formation, and that the bias towards generating new G sites suggests specific DNA repair enzymes may be recruited to these sub-regions within FW1.

## Discussion

In this study, we leveraged deep learning to gain novel insights into SHM, a key process in antibody affinity maturation. We trained multiple deep learning models using a convolutional neural network (CNN) framework to analyze DNA *k*-mer subsequences of various lengths, ranging from 5 to 21 nts, derived from human IGHV germline sequences. Using a high-quality data set containing non-productive B cell repertoire data, the model was tasked to learn two focal aspects of SHM: the frequency of mutation at a given site, and the spectrum of mutations that can arise at this site (substitution). Understanding the propensity of a site to mutate and the underlying substitution biases that ensue can lead to a better understanding of how AID is recruited to and targets the Ig V region, as well as the associated downstream DNA repair mechanisms that follow AID deamination.

We began by developing three models, collectively referred to as DeepSHM, to predict separate tasks for a given *k*-mer: observable mutation frequency; distributed substitution rates; and a combination of both measures (weighted substitution). We found that predicting substitution rates did not substantially depend on the *k*-mer size, while 15-mers were optimal for predicting mutation frequencies (**Figure 2, Table 1**). Additionally, DeepSHM predicted both substitution rates and mutation frequencies more accurately than the widely used S5F targeting model for all *k*-mer sizes we evaluated (*k* = 5, 9, 15 and 21) (**Table 1**). Even though we were able to outperform S5F in representing substitution biases, the correlation between our predictions and empirical data was moderate (∼0.55), suggesting that the processes underlying SHM substitution biases may be more fundamentally random than mutational site targeting alone. Error-prone DNA repair processes downstream of AID are highly complex. For example, while Polη is biased towards making W<u>A</u>>W<u>G</u> mutations (Zhang et al., 2014) and plays a dominant role in generating mutations at A:T sites, many A:T mutations still occur in its absence (Saribasak et al., 2009) that are mediated by other polymerases (Maul et al., 2016). Similarly complex, BER is biased towards transversions but can also repair faithfully, with a further dependence on hotspot mutability (Pérez-Durán et al., 2012). Thus, downstream repair processes may simply be too complex, or genuinely random, to be captured well by a model that depends on sequence context alone.

In order to uncover some of the hidden features learned by DeepSHM, we analyzed the output, or encodings, obtained from the penultimate layer of the network predicting weighted substitution using input 15-mers, and performed t-SNE, a method of dimensionality reduction, to visualize the encodings in two dimensions. The subsequent embedding formed clusters of 15-mers that were distinguished by mutation frequency and middle nucleotide (**Figure 3A, B**). Individual clusters containing a C or G middle nucleotide that were associated with high mutability, assumed to be relevant to AID hotspots, revealed a strong preference for a T base at the +1 position of the top strand AID WR<u>C</u> (W=A/T, R=A/G) hotspot, including for WA<u>C</u> motifs that are not part of a WG<u>C</u>W motif, and similarly, an A base at the −1 position of the bottom strand <u>G</u>YW (Y=C/T) context (**Figure 3C, D**). As an alternative way to identify sequence features, we applied TF-MoDISco (see Methods) to reveal recurrent genomic patterns using importance scores extracted from the model for each 15-mer. This approach confirmed the importance of the T base at the +1 position of WR<u>C</u> (**Figure 4A**) and the A base at the −1 position of the bottom strand <u>G</u>YW hotspot (**Figure 4B**). An early study by Rogozin and Diaz reported the WR<u>C</u>H/D<u>G</u>YW (H=A/C/T, D=A/G/T) to be a good predictor of mutability at C:G bases (Rogozin and Diaz, 2004), but we found WR<u>C</u>T to be a more consistent definition. The authors of the S5F model also supported the WR<u>C</u>H definition since they found their model can capture the higher mutability rate seen at certain WR<u>C</u>A motifs (Yaari et al., 2013), presumably at the A<u>GC</u>A overlapping hotspot.

However, previous hotspot definitions have largely failed to describe targeting beyond the −2 position of the WR<u>C</u> motif. We further identified having a C at the −3 position of WR<u>C</u> or a G at the +3 position of <u>G</u>YW as a strong negative contribution, i.e., as a reduced effect on targeting.

Thus, our results suggest the typical AID hotspot definition might be extended to DWR<u>C</u>T (D=A/G/T). Comparing the mutation frequencies of the individual DWR<u>C</u>T hotspot motifs showed the 3’ T to be more important for AID recognition than the 5’ D alone, however, together they have a synergistic effect that makes mutability between 1.8-fold (for TA<u>C</u>) and 4.7-fold (for TG<u>C</u>) higher (**Figure 5C, Supplementary Table 3**).

We next applied the same t-SNE methodology on the two developed standalone models that separately predicted either the mutation frequency or substitution rates of the 15-mer middle nucleotides. The t-SNE embedding on the independent DeepSHM model predicting only mutation frequency revealed a significant negative correlation between the mutability of a site and the surrounding GC content of the 15-mer (**Figure 7D**). This finding alternatively suggests that highly mutated sites may have evolved to have a richer local AT content. This *in vivo* result is consistent with earlier *in vitro* results that considered AID targeting on artificial substrates (Abdouni et al., 2018).

On the other hand, the t-SNE embedding stemming from the standalone substitution model hinted at plausible associations, both positive and negative, between mutation frequency and certain transition and transversion mutations (**Figure 8B-F**). We next analyzed multiple genes representing different IGHV families containing the largest amounts of mutation data in order to avoid any potential sites with few observable mutations, such as coldspots. We observed a negative correlation between mutability and substitution rates specifically for C>A and G>T transversion mutations (**Figure 8, Supplementary Figure 6**) and, on the other hand, positive correlations for C>T and G>A transitions (**Supplementary Figure 6**). The trend for increased transition mutations at highly mutating AID hotspots mediated by UNG2 had previously been observed in experiments using 3T3 (mouse fibroblast) cells (Pérez-Durán et al., 2012), although the particular bias against C:G>A:T transversions was not apparent. Previous work has also shown that UNG2 is cell-cycle regulated, possibly mediated by FAM72A (Feng et al., 2020), and active primarily during G1 (Sharbeen et al., 2012). Although AID is also primarily active during G1, it may sometimes persist for slightly longer than UNG2 and thus highly targeted sites may avoid BER especially when the mutations occur just before the cell enters S phase, which would lead to fixation of C>T transitions via replication bypass. Alternative polymerases may also be preferentially recruited to some sites. For example, in DT40 (chicken) B-cell lines, the POLD3 subunit of Polymerase delta (Polδ) has been proposed as a specific mechanism for both C>A and G>T mutations (Hirota et al., 2015; Pilzecker and Jacobs, 2019).

Additionally, we investigated two outlier clusters from the substitution model embedding that contained 15-mers having a C and T middle nucleotide that did not group with their respective larger clusters (**Figure 8A**). A closer analysis revealed that the T outlier contained a highly conserved AGYC<u>T</u>GGGGG consensus sequence that was derived from two independent sites located in FW1 from multiple IGHV3 alleles (**Figure 9A, Table 2**). Both outlier clusters also displayed significantly elevated T>G (**Figure 9B, Table 2**) and C>G substitution rates (**Supplementary Figure 7, Table 2**) respectively. In our recent study on G-quadruplexes (G4s) in IGHV genes, we observed the IGHV3 family to form G4s more favorably on the top strand, as measured by their predicted G4 potential using a pre-trained CNN model (Tang and MacCarthy, 2021). Given the strong preference for creating G mutations in these FW1 sub-regions, we evaluated the impact of these mutations on G4 potential. In some cases, the resulting G mutation led to a strong increase in G4 potential, particularly at position 21 (**Figure 9C, Table 2**), whereas for other sites, the effect was mostly negligible (**Table 2**). Notably however, a high A>G substitution rate was also observed at the +3 or +4 positions (**Figure 9A**), which were also associated with increase in G4 potential (**Table 2**). These biased A>G mutations may further be related to previous work that found that a repeated mutation that occurs in one IGHV allele often matches the sequence variant of a different allele (Saini and Hershberg, 2015). Alternatively, these mutations may be related to R-loop initiation, which forms in G-rich non-template DNA, possibly forming in FW1 of these IGHV3 genes. Studies have found that reducing G-density in mammalian Ig Switch regions compromises class-switch recombination efficiency and R-loops from forming (Roy et al., 2008; Zhang et al., 2014). The high rate of T>G and C>G transversions also suggests that particular repair enzymes may be recruited to these sub-regions during SHM.

### Limitations of the study

In principle, a wider range of k-mers, as well as a greater variety of neural network architectures, might have been considered for this study. However, since the tuning of each model takes a substantial amount of computational resources and time, we considered a reduced number of models. Additionally, we limited this study to consider data only for human, the species for which we had high quality (UMI barcoded) data in high abundance, although the approach could be extended to other species such as mouse in future work.

## Acknowledgements

We would like to extend our gratitude to Sergio Roa Gómez for his comments and suggestions.

## Author contributions

C.T., A.K., and T.M. conceived the idea, analyzed the results, and wrote the manuscript. C.T. and A.K. developed the model and performed computational analysis. All authors contributed to the article and approved the submitted version.

## STAR Methods

### Resource availability

#### Lead contact

Further information and requests for resources and reagents should be directed to and will be fulfilled by the lead contact, Thomas MacCarthy.

## Materials availability

This study did not generate new unique reagents.

## Data and code availability

Data used for this research was published previously by Tang et al, 2020. A custom Python package developed for this project is available at https://gitlab.com/maccarthyslab/deepshm.

## Methods details

### Generating *k*-mer data

Germline IGHV reference sequences were downloaded from the international ImMunoGeneTics information system (IMGT) website (Lefranc, 2001). The leader portion of each reference sequence was also extracted if available. To generate the *k*-mers of a given germline sequence, ±⌊*k*/2⌋ nt sequences were extracted from the start of the V exon, where *k* is the length of the subsequence, and ⌊*k*/2⌋ represents the greatest integer less than or equal to *k*/2. This process was continued, moving 1 nt at a time, until the end of the exon was reached. Next, all *k*-mers were converted to their respective one-hot encodings. A one-hot encoding is a transformation of a DNA sequence using a 2-D matrix containing only zeros and ones, where each row represents one of the four ordered DNA bases and each column is an individual site in the sequence. For each column, a “1” is filled in the row that matches the nucleotide of that site and a “0” in the remaining unmatched rows (**Figure 1**).

### Calculating mutation frequencies, substitution rates, and weighted substitutions of *k*-mers

Using a high-quality data set previously published by us (Tang et al., 2020), we calculated the mutation frequencies of every *k*-mer in a germline sequence as the number of observed mutations at each site (corresponding to a single *k*-mer), divided by the total number of sequences the germline IGHV allele contained. The substitution rate of each *k*-mer was computed as the number of times the middle nucleotide mutated from the germline nucleotide to the other four DNA bases, divided by the total number of overall mutations. Note that a zero was recorded in the instance the mutated base was the same as the germline context. Lastly, the weighted substitution of a *k*-mer was simply calculated as the observed mutation frequency multiplied by the substitution rate vector.

### CNN architecture and model optimization

We implemented a convolutional neural network (CNN) to analyze the *k*-mer input data. Three separate architectures were used to predict different SHM outcomes: mutation frequency, substitution rate, and weighted substitution (see above). Although the hyperparameters that were ultimately selected varied from model-to-model, all CNNs followed the same general architecture, which consisted of one convolution layer, followed by two fully connected layers (**Figure 1**). Additional parameters, such as dropout and batch normalization, were optimized by generating 100 separate models with randomly selected hyperparameters for each *k*-mer and corresponding model architecture we generated. The range of values for all parameters and hyperparameters that were tested for each architecture and output type are specified in **Supplementary Table 1**.

Next, we utilized 4-fold cross-validation to evaluate the performance of the model on unseen (test) data. In total, there are seven IGHV families (IGHV1-7), where each IGHV family consists of genes that share a high percentage of sequence similarity (Lefranc, 2001). The *k*-mers derived from the three largest IGHV families, IGHV1, IGHV3, and IGHV4, formed three separate groups, and the *k*-mers belonging to the remaining 4 smaller IGHV families constituted the final group in order to create a data set comparable in size with the other groups. Thus, we separated the data by their respective IGHV family to reduce the chances of model overfitting, since it is likely that *k*-mers from the same IGHV family will be similar even if they come from different genes and, therefore, bias the results if they appear in both training and test sets. In every cross-validation fold, three of the data groups were used as training set, and the fourth used as test set. We also evaluated the model performance, for each fold, by calculating the Pearson correlation(s) between the predicted mutation frequency and/or substitution rate of the test set *k*-mers and the equivalent output type of the empirical data. The average correlation across the 4 validation folds was reported for the model, as in **Figure 2**.

As an additional step, we wrote a custom, universal Python script (available at https://gitlab.com/maccarthyslab/deepshm) to automatically generate the CNN architecture, parameters, and hyperparameters of each model, regardless of the output specified, to ensure that all models were constructed in a consistent manner. All CNNs were generated using the built-in Keras API in Tensorflow 2.4.1 and trained on GPU processors using three Nvidia GeForce RTX 2080 graphics cards.

### Inferring an S5F targeting model

In order to ensure a fair comparison between S5F values and our deep learning predictions, we used the SHazaM R package (Yaari et al., 2013) to create an S5F targeting model, which provides analogous 5-mer mutability and substitution scores based on the same data set we used to train our CNN models with. We specified the S5F targeting model to count both silent and replacement mutations (“rs” parameter) since the mutation data we used was derived from non-functionally rearranged VDJ coding sequences (i.e. in the absence of selection) and with each sequence being clonally independent (Tang et al., 2020). Multiple mutations were handled specifying the “independent” parameter, which treats each mutation independently. Default values were used for all other parameters.

## Quantification and statistical analysis

### Neural network encodings analysis

The output (encoding) of the penultimate layer of the CNN model was used as a way to explain the SHM patterns learned by the model. To generate the encodings from this layer, we removed the last layer of the CNN while keeping the remaining layers intact. Next, we processed the *k*-mers through the truncated model to retrieve the ensuing output values. We then applied t-distributed stochastic neighbor embedding (t-SNE) in Python on these multidimensional encodings to visually represent the resulting embedding in two dimensions.

### Cluster identification

We implemented k-means clustering to identify clusters within the t-SNE embedding of the weighted substitution model (**Supplementary Figure 1**). We separated all *k*-mers sharing the same middle nucleotide and then applied k-means clustering independently on each group to facilitate the clustering process. All clustering assignments were performed using the *kmeans* function in R. For each middle nucleotide, we specified the algorithm to identify 5 distinct clusters and subsequently inspected the clusters to ensure a proper separation between clusters of distinct mutabilities occurred. In the case of G and T nucleotides, there were resulting clusters (one for each nucleotide) containing hot and cold sequences (i.e one “cold” and one “hot” subcluster per cluster), so we manually split each of these clusters into two distinct (“cold” and “hot”) clusters to reduce the disparity in mutation frequencies.

### Identifying recurring genomic patterns using TF-MoDISco

We applied TF-MoDISco (Shrikumar et al., 2018), a machine learning interpretability method, to identify recurring motifs our model detected in the 15-mer data. From the data, we isolated four groups of 15-mers based on the middle nucleotide (A, C, G, or T) of the 15-mer, with the additional condition that the middle nucleotide conformed to WR<u>C</u> or <u>G</u>YW AID hotspots, or W<u>A</u> or <u>T</u>W Polη hotspots, respectively. TF-MoDISco requires importance scores to be used as input, which can be generated by utilizing one of several attribution methods. Here we generated the importance scores for each group by applying Integrated Gradients (Sundararajan et al., 2017) to the most accurate 15-mer mutation frequency model. Using the resulting importance scores, we then ran TF-MoDISco for all groups separately, still subject to the hotspot constraint, and requiring each of the identified patterns to be associated with at least 20 input sub-sequences (or “sequelets”).

## Key resources table

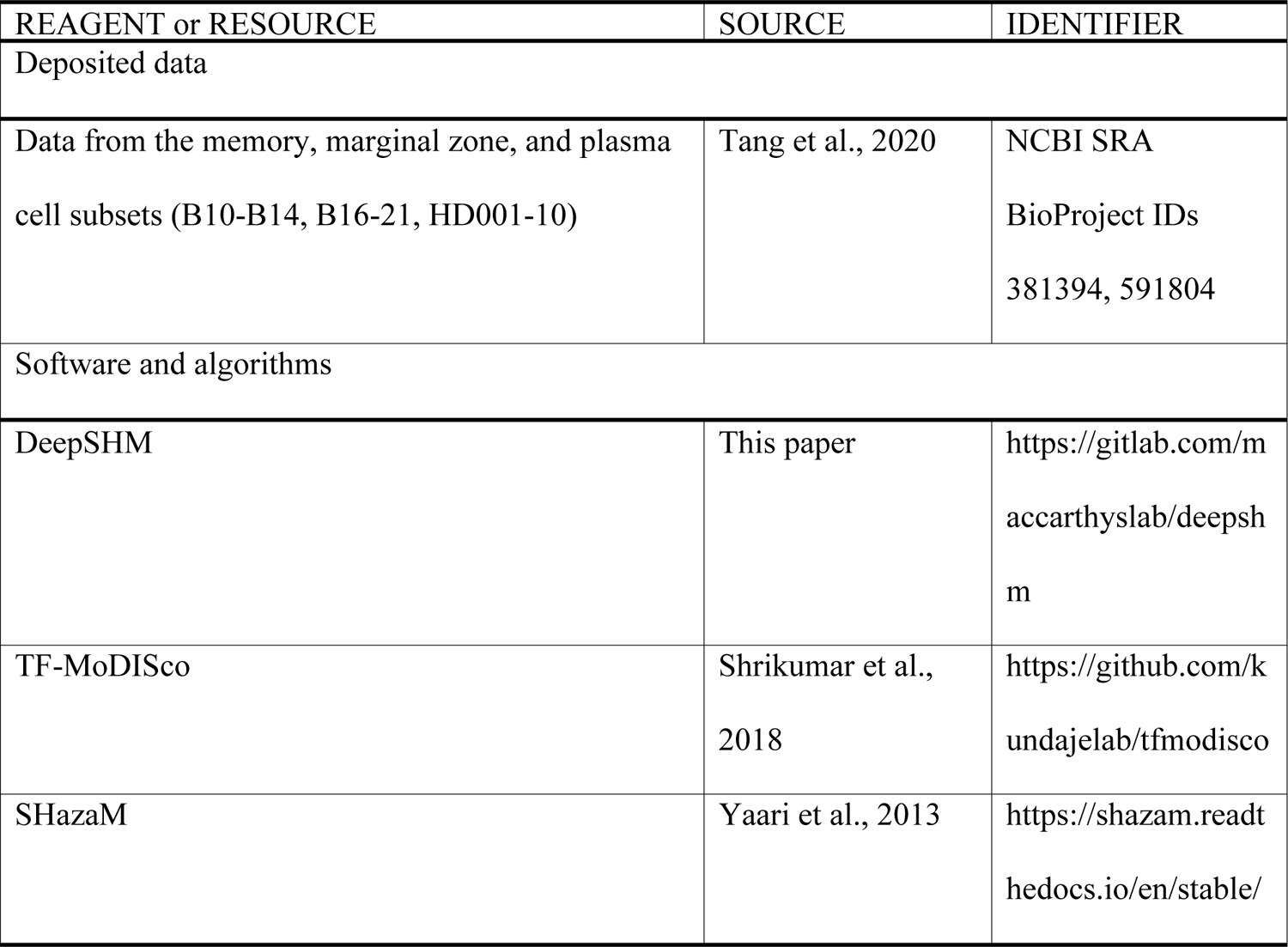

