## Supplemental Materials for "Deep learning model of somatic hypermutation reveals importance of sequence context beyond targeting of AID and Polη hotspots"

**Supplementary Table 1. List of parameters and hyperparameters tested.**

| Parameter | Range of tested values |
| --- | --- |
| Input *k* -mer length (k) | 5, 9, 15, 21 |
| Task | Substitution, mutation frequency, weighted substitution |
| Number of channels (convolution layer) | 64, 128, 256 |
| Kernel size | [4, 2] to [4, k] |
| Average pooling | N/A, [2, 1] |
| Dropout rate (convolution layer) | 0.0, 0.1, 0.2, 0.3, 0.4, 0.5 |
| Number of channels (1st fully connected layer) | 16, 32, 64, 128, 256 |
| Number of channels (2nd fully connected layer) | 4, 8, 16, 32 |
| Dropout rate (fully connected layers) | 0.0, 0.1, 0.2, 0.3, 0.4, 0.5 |
| Number of epochs | 100, 150, 200 |
| Learning rate | 2-11, 2-12, 2-13, 2-14, 2-15 |
| Mini-batch size | 8, 16, 32 |

**Supplementary Table 2. Cluster analsyis.**

| Cluster | n | Middle nucleotide | Avg. mutation frequency | Consensus |
|---|---|---|---|---|
| 1 | 899 | A | 0.14 ± 0.10 | 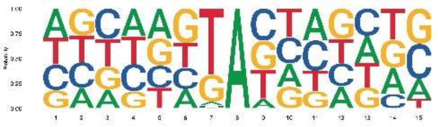 |
| 2 | 715 | A | 0.13 ± 0.06 | 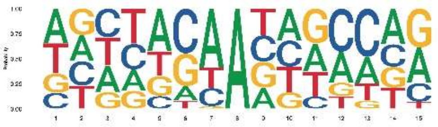 |
| 3 | 1208 | A | 0.06 ± 0.04 | 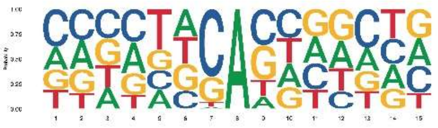 |
| 4 | 782 | A | 0.03 ± 0.02 | 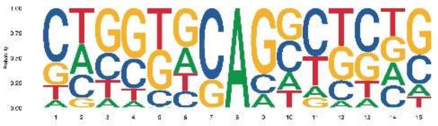 |
| 5 | 1823 | A | 0.07 ± 0.04 | 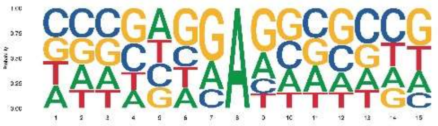 |
| 6 | 1606 | C | 0.05 ± 0.04 | 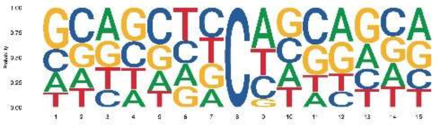 |
| 7 | 1530 | C | 0.11 ± 0.06 | 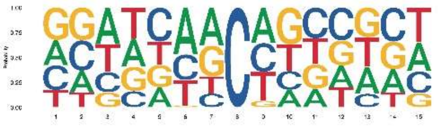 |
| 8 | 752 | C | 0.02 ± 0.03 | 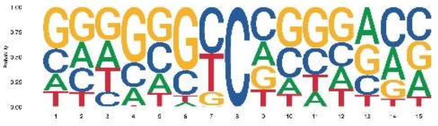 |
| 9 | 1415 | C | 0.02 ± 0.02 | 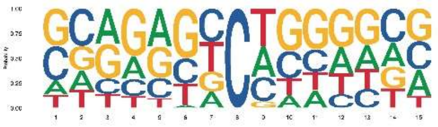 |
| 10 | 538 | C | 0.24 ± 0.08 | 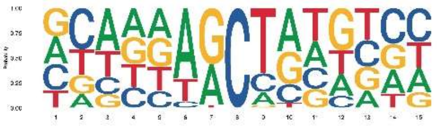 |
| 11 | 988 | G | 0.03 ± 0.02 | 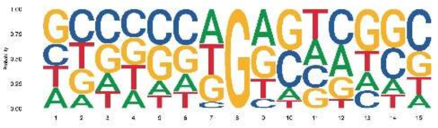 |

| | | | | |
|---|---|---|---|---|
| 12 | 602 | G | 0.02 ± 0.02 | 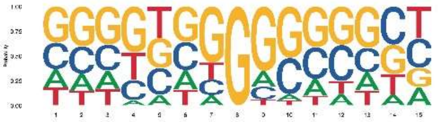 |
| 13 | 2122 | G | 0.05 ± 0.04 | 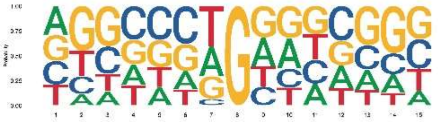 |
| 14 | 946 | G | 0.02 ± 0.02 | 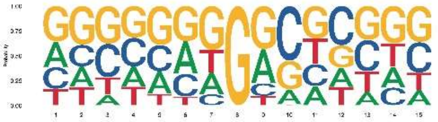 |
| 15 | 1907 | G | 0.13 ± 0.09 | 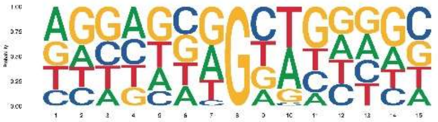 |
| 16 | 283 | G | 0.37 ± 0.10 | 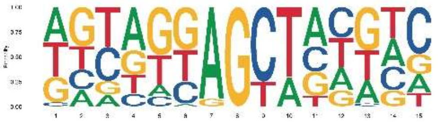 |
| 17 | 1220 | T | 0.04 ± 0.04 | 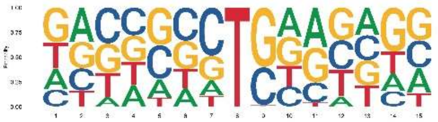 |
| 18 | 1284 | T | 0.06 ± 0.04 | 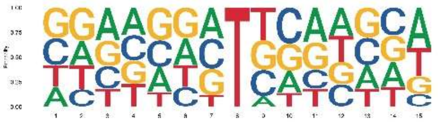 |
| 19 | 575 | T | 0.03 ± 0.03 | 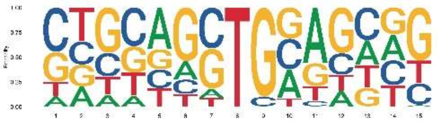 |
| 20 | 878 | T | 0.13 ± 0.07 | 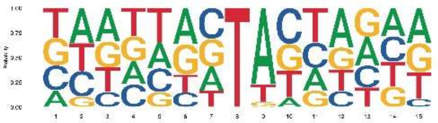 |
| 21 | 684 | T | 0.02 ± 0.02 | 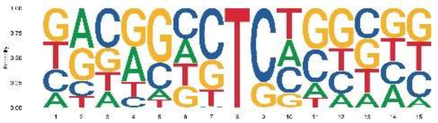 |
| 22 | 101 | T | 0.07 ± 0.04 | 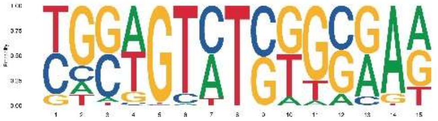 |

**Supplementary Table 3. Pairwise comparison of extended AID hotspot mutabilities.**

| Extended hotspot | n | Avg. mutation frequency | DAGCT | DTGCT | DAACT | DTACT | CAACT | CAGCT | CTACT | DAACV | DTACV | CTGCT | DAGCV | CTACV | CAACV | CAGCV | DTGCV | CTGCV |
|---|---|---|---|---|---|---|---|---|---|---|---|---|---|---|---|---|---|---|
| DAGCT | 236 | 0.32 ± 0.12 | 9.48E-01 | 3.72E-02 | 1.56E-04 | 5.25E-05 | 1.76E-06 | 3.93E-23 | 1.88E-25 | 6.64E-37 | 3.08E-39 | 7.32E-22 | 2.18E-52 | 5.50E-35 | 3.47E-23 | 1.27E-61 | 8.06E-71 | 1.75E-85 |
| DTGCT | 111 | 0.28 ± 0.10 | 1.00E+00 | 9.48E-01 | 8.69E-02 | 4.00E-02 | 1.12E-04 | 2.98E-10 | 4.54E-15 | 5.26E-22 | 2.34E-22 | 1.65E-16 | 4.87E-28 | 4.14E-23 | 5.09E-17 | 8.70E-35 | 1.17E-40 | 4.84E-51 |
| DAACT | 116 | 0.27 ± 0.13 | 1.00E+00 | 1.00E+00 | 9.48E-01 | 9.06E-01 | 2.53E-02 | 1.35E-06 | 7.06E-09 | 2.68E-17 | 8.80E-17 | 5.21E-12 | 9.30E-23 | 2.61E-19 | 4.09E-15 | 3.04E-31 | 7.22E-38 | 5.46E-52 |
| DTACT | 80 | 0.25 ± 0.09 | 1.00E+00 | 1.00E+00 | 1.00E+00 | 9.48E-01 | 2.23E-02 | 6.78E-05 | 3.85E-08 | 7.62E-14 | 1.12E-13 | 1.12E-11 | 8.51E-18 | 2.43E-16 | 2.07E-13 | 3.15E-24 | 3.44E-29 | 1.16E-39 |
| CAACT | 27 | 0.20 ± 0.08 | 1.00E+00 | 1.00E+00 | 1.00E+00 | 1.00E+00 | 9.48E-01 | 6.28E-01 | 9.76E-02 | 3.06E-03 | 1.93E-03 | 2.43E-04 | 1.98E-04 | 1.52E-04 | 1.48E-05 | 8.69E-07 | 3.34E-09 | 1.29E-14 |
| CAGCT | 298 | 0.20 ± 0.12 | 1.00E+00 | 1.00E+00 | 1.00E+00 | 1.00E+00 | 1.00E+00 | 9.48E-01 | 6.28E-01 | 3.45E-05 | 3.22E-06 | 3.73E-03 | 7.18E-11 | 1.11E-07 | 1.36E-06 | 5.32E-20 | 4.82E-31 | 1.73E-59 |
| CTACT | 131 | 0.19 ± 0.08 | 1.00E+00 | 1.00E+00 | 1.00E+00 | 1.00E+00 | 1.00E+00 | 1.00E+00 | 9.48E-01 | 4.76E-05 | 9.60E-05 | 2.60E-03 | 6.65E-09 | 9.76E-09 | 3.44E-08 | 2.87E-18 | 3.13E-26 | 2.77E-49 |
| DAACV | 173 | 0.16 ± 0.08 | 1.00E+00 | 1.00E+00 | 1.00E+00 | 1.00E+00 | 1.00E+00 | 1.00E+00 | 1.00E+00 | 9.48E-01 | 4.52E-01 | 1.00E+00 | 3.37E-02 | 6.99E-02 | 5.39E-03 | 7.75E-07 | 2.90E-14 | 2.34E-41 |
| DTACV | 215 | 0.15 ± 0.10 | 1.00E+00 | 1.00E+00 | 1.00E+00 | 1.00E+00 | 1.00E+00 | 1.00E+00 | 1.00E+00 | 1.00E+00 | 9.48E-01 | 1.00E+00 | 3.00E-01 | 3.92E-01 | 6.39E-02 | 1.98E-04 | 5.99E-11 | 5.15E-32 |
| CTGCT | 56 | 0.15 ± 0.07 | 1.00E+00 | 1.00E+00 | 1.00E+00 | 1.00E+00 | 1.00E+00 | 1.00E+00 | 1.00E+00 | 8.20E-01 | 8.45E-01 | 9.48E-01 | 1.47E-01 | 6.99E-02 | 1.32E-02 | 2.28E-04 | 6.26E-08 | 5.24E-19 |
| DAGCV | 343 | 0.14 ± 0.01 | 1.00E+00 | 1.00E+00 | 1.00E+00 | 1.00E+00 | 1.00E+00 | 1.00E+00 | 1.00E+00 | 1.00E+00 | 1.00E+00 | 1.00E+00 | 9.48E-01 | 9.92E-01 | 2.66E-01 | 2.88E-03 | 8.79E-11 | 8.56E-39 |
| CTACV | 124 | 0.14 ± 0.08 | 1.00E+00 | 1.00E+00 | 1.00E+00 | 1.00E+00 | 1.00E+00 | 1.00E+00 | 1.00E+00 | 1.00E+00 | 1.00E+00 | 1.00E+00 | 9.48E-01 | 9.48E-01 | 2.28E-01 | 1.22E-02 | 3.49E-07 | 5.99E-27 |
| CAACV | 68 | 0.13 ± 0.10 | 1.00E+00 | 1.00E+00 | 1.00E+00 | 1.00E+00 | 1.00E+00 | 1.00E+00 | 1.00E+00 | 1.00E+00 | 1.00E+00 | 1.00E+00 | 1.00E+00 | 1.00E+00 | 9.48E-01 | 5.94E-01 | 4.63E-03 | 4.81E-12 |
| CAGCV | 330 | 0.12 ± 0.09 | 1.00E+00 | 1.00E+00 | 1.00E+00 | 1.00E+00 | 1.00E+00 | 1.00E+00 | 1.00E+00 | 1.00E+00 | 1.00E+00 | 1.00E+00 | 1.00E+00 | 1.00E+00 | 1.00E+00 | 9.48E-01 | 2.85E-04 | 2.01E-27 |
| DTGCV | 333 | 0.10 ± 0.08 | 1.00E+00 | 1.00E+00 | 1.00E+00 | 1.00E+00 | 1.00E+00 | 1.00E+00 | 1.00E+00 | 1.00E+00 | 1.00E+00 | 1.00E+00 | 1.00E+00 | 1.00E+00 | 1.00E+00 | 1.00E+00 | 9.48E-01 | 7.34E-11 |
| CTGCV | 333 | 0.06 ± 0.05 | 1.00E+00 | 1.00E+00 | 1.00E+00 | 1.00E+00 | 1.00E+00 | 1.00E+00 | 1.00E+00 | 1.00E+00 | 1.00E+00 | 1.00E+00 | 1.00E+00 | 1.00E+00 | 1.00E+00 | 1.00E+00 | 1.00E+00 | 9.48E-01 |

P-values from one-sided Mann-Whitney U tests, where the hotspot in the row is tested for significantly higher mutability than the hotspot in the column.

P-values are adjusted for multiple comparisons (Benjamini-Hochberg corrected).

Significant p-values are highlighted in red.

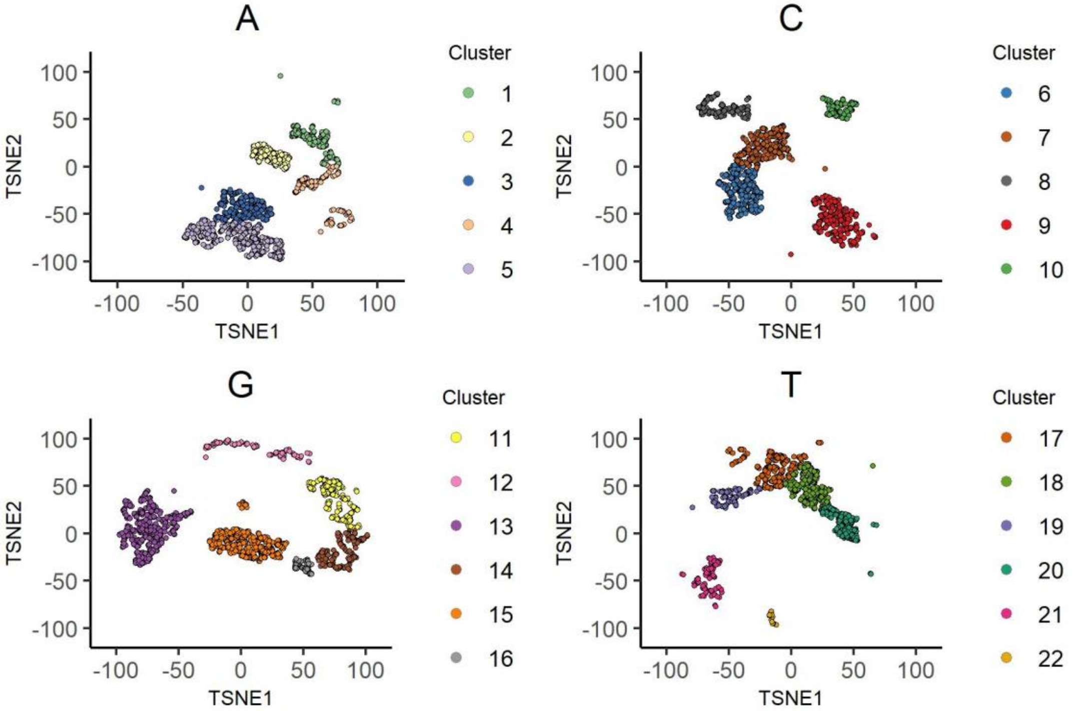
A
C
G
T
Cluster
TSNE1
TSNE2
100
50
0
-50
-100

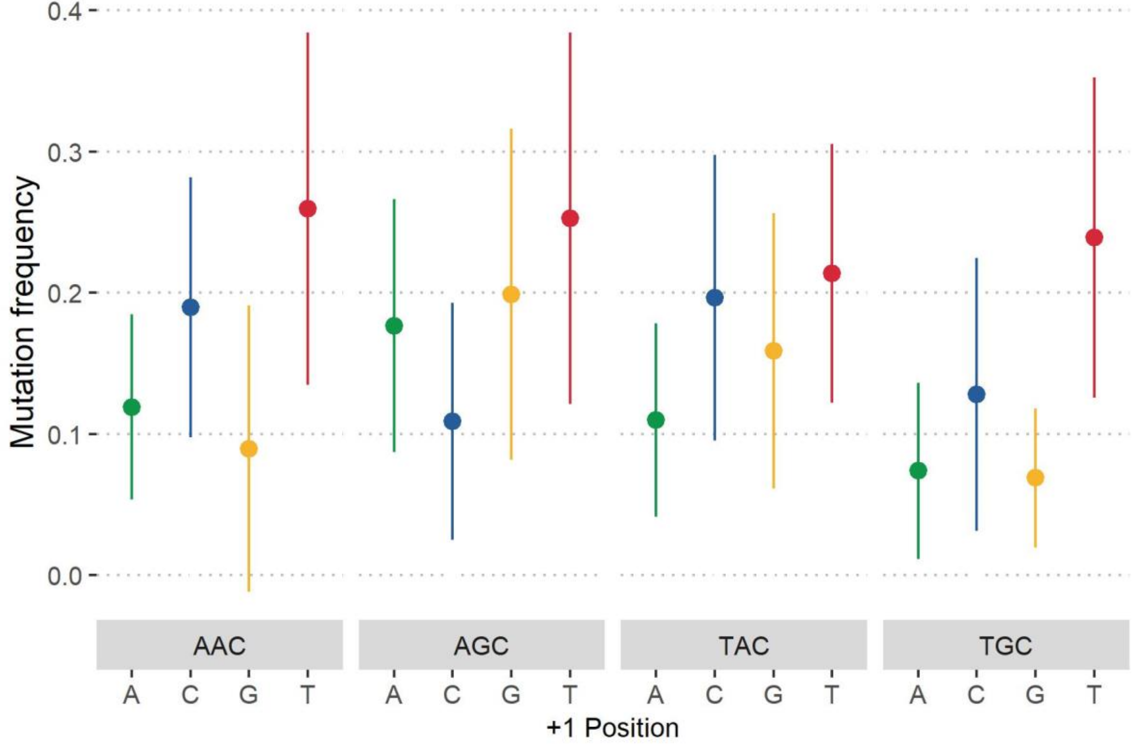

Mutation frequency
0.4
0.3
0.2
0.1
0.0
AAC
AGC
TAC
TGC
A
C
G
T
+1 Position

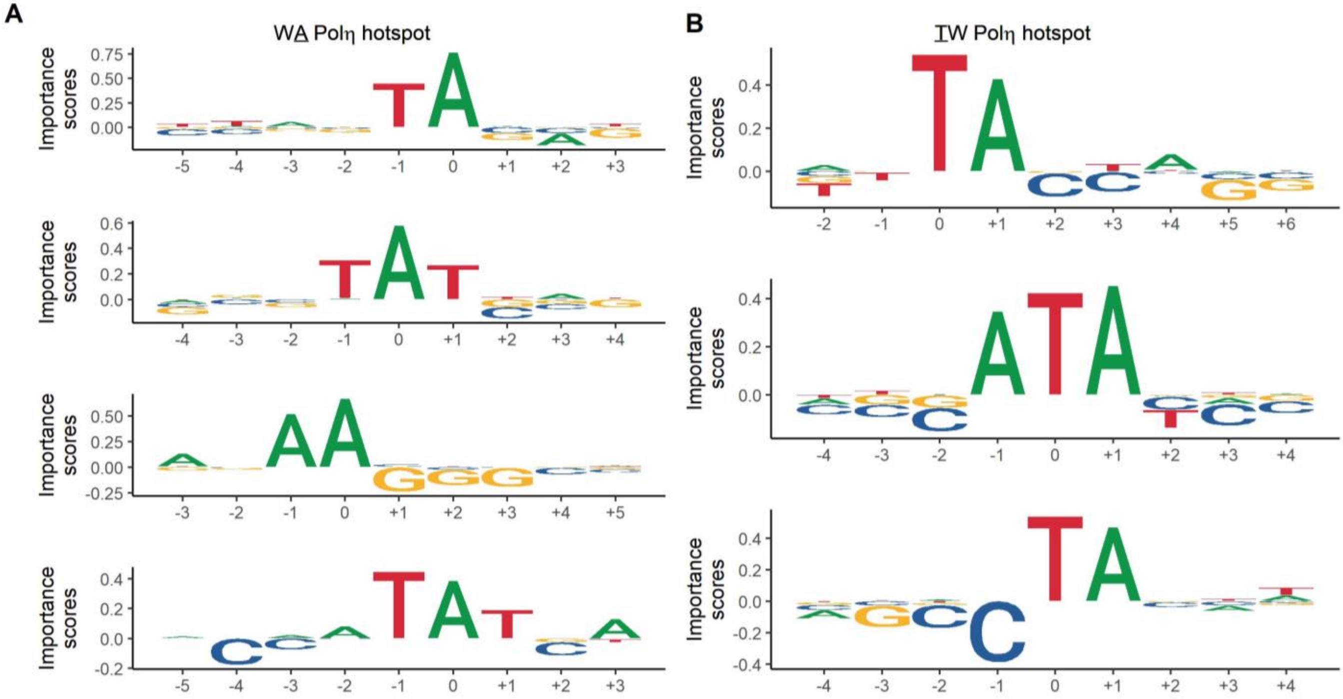

A
WA Polη hotspot
B
TW Polη hotspot
Importance scores

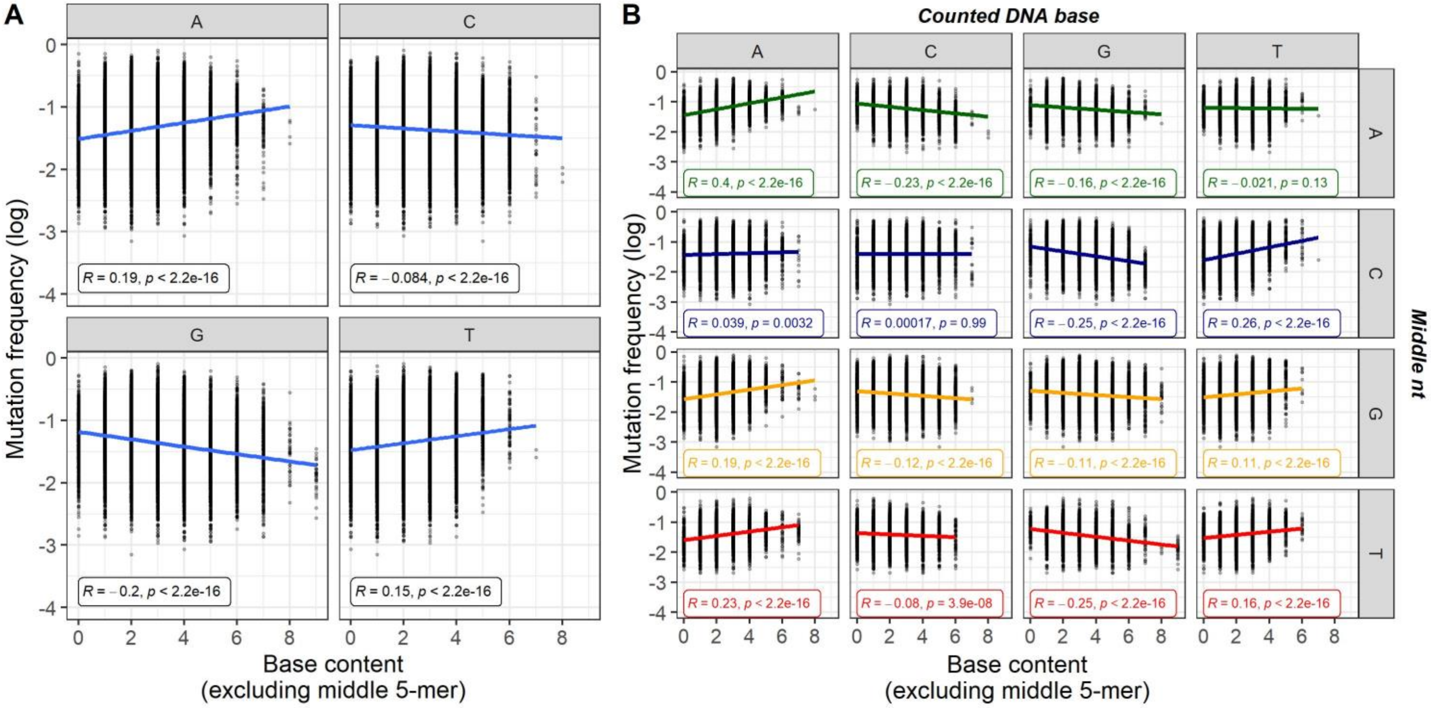

A
B
Counted DNA base
Middle nt
A
C
G
T
Mutation frequency (log)
Base content
(excluding middle 5-mer)
R = 0.19, p < 2.2e-16
R = −0.084, p < 2.2e-16
R = −0.2, p < 2.2e-16
R = 0.15, p < 2.2e-16
R = 0.4, p < 2.2e-16
R = −0.23, p < 2.2e-16
R = −0.16, p < 2.2e-16
R = −0.021, p = 0.13
R = 0.039, p = 0.0032
R = 0.00017, p = 0.99
R = −0.25, p < 2.2e-16
R = 0.26, p < 2.2e-16
R = 0.19, p < 2.2e-16
R = −0.12, p < 2.2e-16
R = −0.11, p < 2.2e-16
R = 0.11, p < 2.2e-16
R = 0.23, p < 2.2e-16
R = −0.08, p = 3.9e-08
R = −0.25, p < 2.2e-16
R = 0.16, p < 2.2e-16

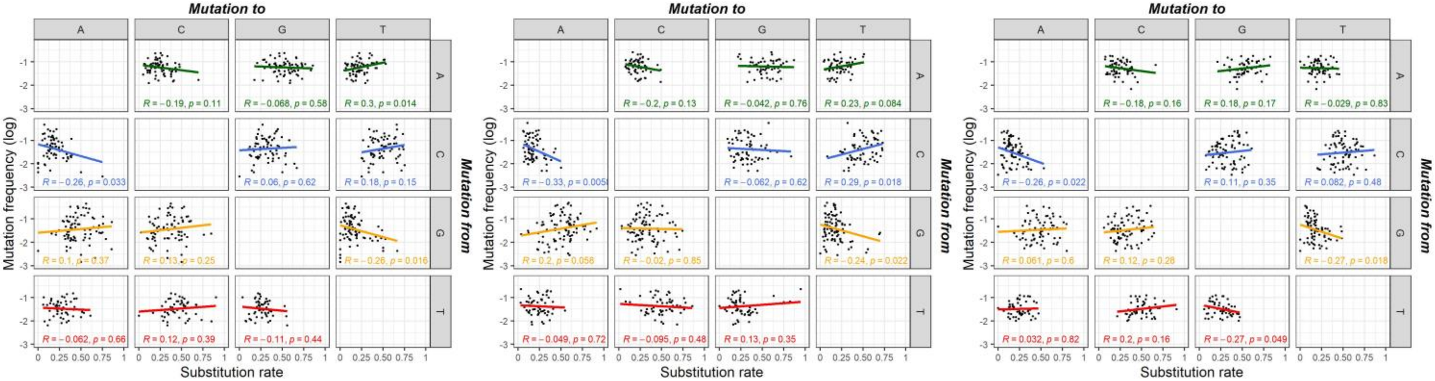

IGHV1-18*01 IGHV3-23*01 IGHV4-34*01

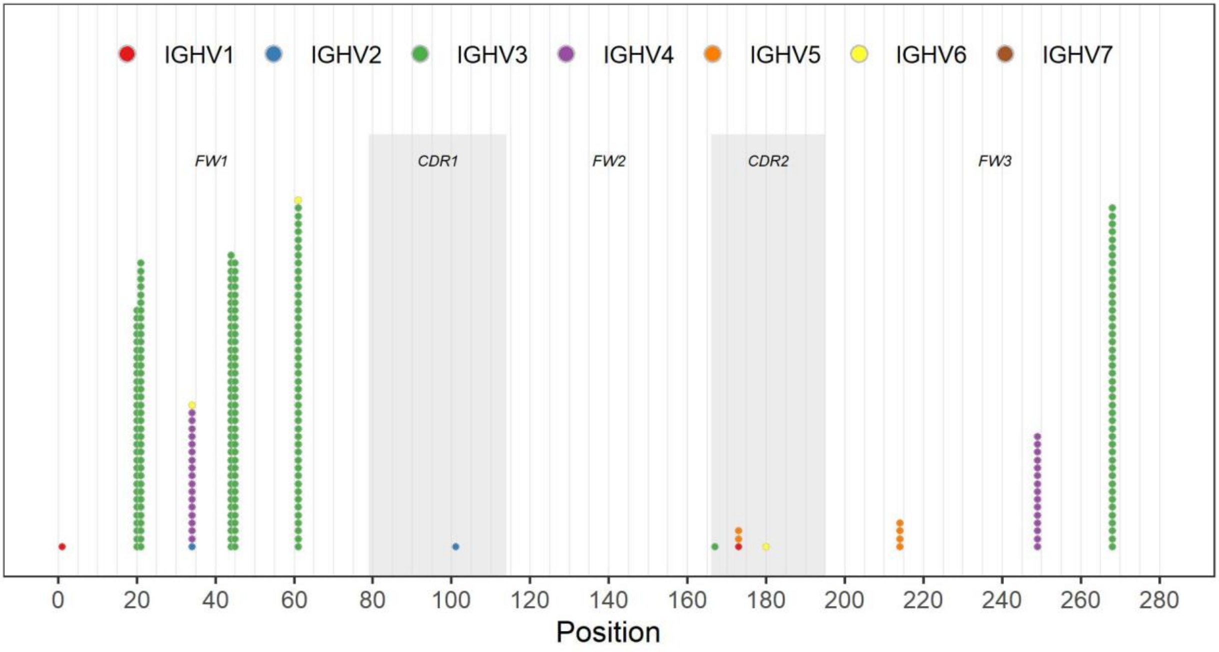

IGHV1
IGHV2
IGHV3
IGHV4
IGHV5
IGHV6
IGHV7
FW1
CDR1
FW2
CDR2
FW3
0
20
40
60
80
100
120
140
160
180
200
220
240
260
280
Position

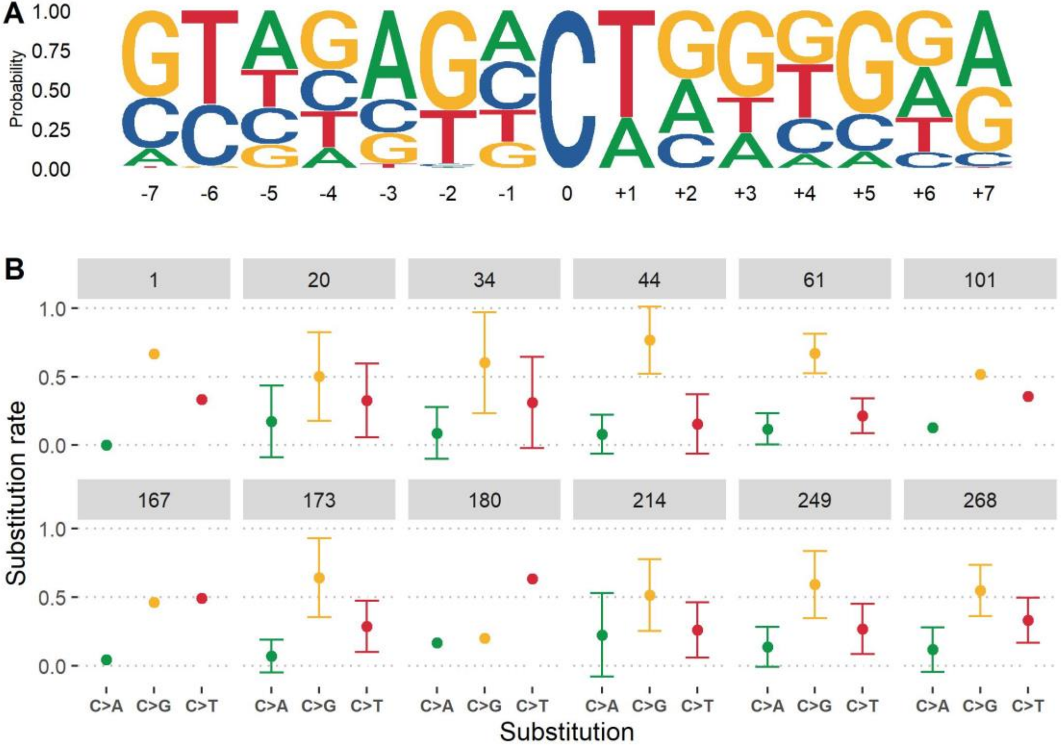

A
Probability
1.00
0.75
0.50
0.25
0.00
-7
-6
-5
-4
-3
-2
-1
0
+1
+2
+3
+4
+5
+6
+7
B
1
20
34
44
61
101
167
173
180
214
249
268
1.0
0.5
0.0
Substitution rate
C>A
C>G
C>T
Substitution

**Supporting Information**

**Supplementary Table 1. List of parameters and hyperparameters tested.**

**Supplementary Table 2. Cluster analysis.**

**Supplementary Table 3. Pairwise comparison of extended AID hotspot mutabilities.**

**Supplementary Figure 1. Clustering performed on the t-SNE embedding of the DeepSHM weighted substitution model**. Cluster plots are separated by middle nucleotide of each 15-mer. Individual clusters are distinguished by different colors.

**Supplementary Figure 2. Mutability assessment of the +1 position of WRCN.** Mean mutation frequencies are shown for all individual WRCN motifs (dots), where the DNA base on the x-axis represents the +1 position of the core WRC motif labeled in gray above. Lines extend +/- 1 standard deviation from the mean.

**Supplementary Figure 3. Recurrent motifs identified by TF-MoDISco.** TF-MoDISco results using the Integrated Gradients as base-level importance scores of 15-mers whose middle nucleotide conformed to a **(A)** W<u>A</u> or **(B)** <u>T</u>W Polη hotspot motif.

**Supplementary Figure 4. Relationship between mutation frequency and individual base content. (A)** Pearson correlations between the individual base count (x-axis) for each 15-mer, excluding the middle 5-mer count, whose middle nucleotide matches the DNA base labelled in gray, and its corresponding mutation frequency (y-axis; log10). **(B)** Same comparison as **(A)** but grouped by 15-mers containing the same middle nucleotide (nt) indicated by the row.

**Supplementary Figure 5. Comparison of mutation frequency and substitution rates in multiple IGHV genes.** Plots for IGHV1-18*01, IGHV3-23*01, and IGHV4-34*01 compare the observed mutation frequency and rates of substitution for each individual mutation type. The DNA base in the row indicates the base subjected to mutation, and the resulting base by the column. Pearson correlations are calculated for each plot separately.

**Supplementary Figure 6. Dotplot of the outlier clusters in the DeepSHM substitution model.** Each dot represents a single 15-mer from either of the outlier clusters identified in **Figure 8A** (red and blue clusters, right) and is colored according to its respective IGHV family. The IMGT position of each 15-mer middle nucleotide is indicated by the x-axis. CDR boundaries are shown in gray.

**Supplementary Figure 7. Evaluation of the C outlier cluster in the DeepSHM substitution model. (A)** Sequence logo representation of the 15-mers appearing in the C outlier cluster in **Figure 9A** (right-hand side, blue dots). **(B)** Substitution rates of C>A, C>G, and C>T separated by the IMGT position indicated in gray. Bars represent ±1 standard deviation.
